# A Small Molecule Fluorogenic Probe for the Detection of Sphingosine in Living Cells

**DOI:** 10.1101/2020.06.27.175661

**Authors:** Andrew K. Rudd, Neel Mittal, Esther W. Lim, Christian M. Metallo, Neal K. Devaraj

**Affiliations:** Department of Chemistry and Biochemistry, University of California San Diego, La Jolla, CA, USA; Department of Bio-engineering, University of California San Diego, La Jolla, CA, USA

## Abstract

The single-chained sphingolipid sphingosine is an essential structural lipid and signaling molecule. Abnormal sphingosine metabolism is observed in several diseases, including cancer, diabetes, and Alzheimer’s. Despite its biological importance, there are a lack of tools for detecting sphingosine in living cells. This is likely due to the broader challenge of developing highly selective and live-cell compatible affinity probes for hydrophobic lipid species. In this work, we have developed a small molecule fluorescent turn-on probe for labeling sphingosine in living cells. This probe utilizes a selective reaction between sphingosine and salicylaldehyde esters to fluorescently label sphingosine molecules. We demonstrate that this probe exhibits a dose-dependent response to sphingosine and is able to detect endogenous pools of sphingosine. Using our probe, we successfully detected sphingosine accumulation in live Niemann-Pick type C1 (NPC1) patient cells, a lipid transport disorder in which increased sphingosine mediates disease progression. This work provides a simple and accessible method for the detection of sphingosine and should facilitate study of this critical signaling lipid in biology and disease.

---

Sphingolipids are a diverse class of lipids defined by their long-chain amino alcohol backbones. In eukaryotes, these lipids play essential roles in both membrane structure and cell signaling pathways.^1^ Several sphingolipid species such as ceramides, sphingomyelin, glucosylceramide, sphingosine, and sphingosine-1-phosphate have emerged as integral signaling molecules in cell proliferation,^2^ apoptosis,^3^ migration,^4^ inflammation,^5^ and intracellular trafficking.^6^ Due to their central role in cellular function, disruption of sphingolipid metabolism can have devastating biological effects. Altered sphingolipid levels are associated with several diseases such as diabetes,^7^ cancer,^8^ Alzheimer’s,^9^ and lysosomal storage diseases.^10,11^ Because of their biological and clinical importance, there is tremendous interest in detecting and quantifying sphingolipids in cells and biological samples. Mass spectrometry-based methods are the current standard for detecting cellular sphingolipids.^12,13^ These techniques have made it possible to accurately measure the abundance of specific sphingolipids in a wide range of samples but often require sophisticated instrumentation and preclude the nondestructive analysis of lipids in living cells. Recently, methods have been developed for the live-cell imaging of sphingomyelin using fluorescent protein fusions of the sphingomyelin binding protein lysenin, which binds specifically to sphingomyelin, however the extension of this approach to other sphingolipids has been limited.^14,15^ Sphingosine (Sph) and sphinganine (Spa) are single-chained sphingolipids which not only serve as the backbones of more complex sphingolipids but also have important biological signaling activity.^16,17^ They play essential roles in cellular function and diseases such as cancer^18^ and Niemann-Pick disease type C (NPC).^10^ Unfortunately, dissecting the exact function and behavior of sphingosines in biology and disease has been difficult due to the lack of effective techniques for their imaging and detection in live cells. The development of such tools is critical for the biological understanding, diagnosis, and treatment of sphingolipid-associated diseases.

The live-cell detection of lipids is hampered by the general hydrophobicity of lipid species and their incredible functional diversity. A unique characteristic of sphingosine and sphinganine, compared to other lipids, is their terminal 1,2-amino alcohol functionality. Recent studies have shown that fatty acid salicylaldehyde esters and 1,2-amino alcohols, such as those found in Sph and Spa, can react selectively under biological conditions.^19,20^ This method results in the transfer of the acyl group from salicylaldehyde to the sphingosine amine, yielding a stable amide linkage. We hypothesized that a similar approach could be taken in the design of a fluorescent probe for Sph and Spa. By synthesizing a fluorophore ester of salicylaldehyde, we hoped we could covalently attach the fluorophore to endogenous Sph and Spa. To aid detection, we wanted the reaction between our probe and the target sphingolipids to be fluorogenic. We hypothesized that this could be accomplished by functionalizing the salicylaldehyde auxiliary with a fluorescence quencher, diminishing the fluorescence of the fluorophore ester. Upon reaction with Sph or Spa, the fluorophore would be transferred from the salicylaldehyde scaffold to the sphingolipid base and become unquenched due to its decreased proximity to the quencher^21^ (Figure 1A). In mammalian cells, Sph is 10-fold more abundant than Spa, and would be the predominant species detected by such a probe.^22^ The envisioned approach would enable the fluorescent detection of Sph in living cells.

**Figure 1.**
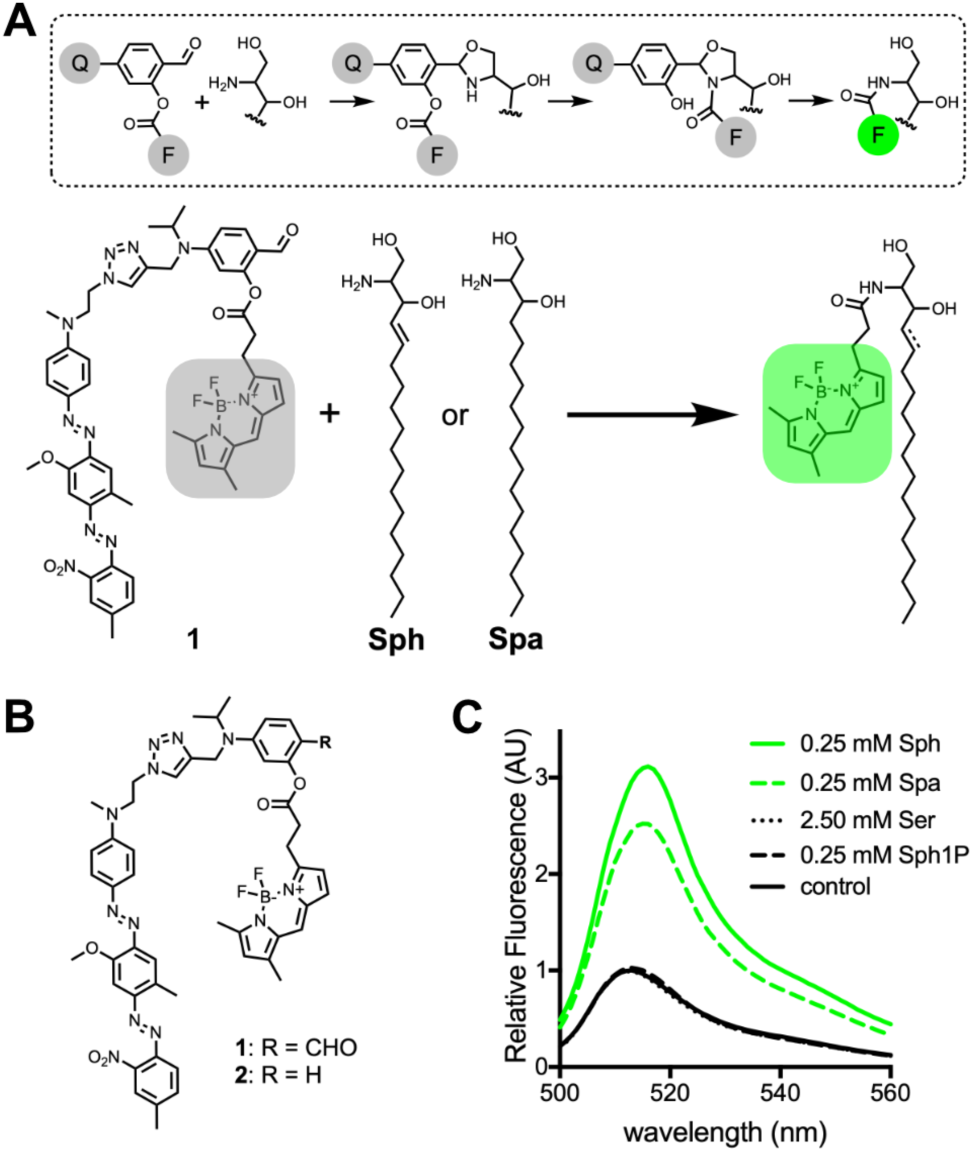
(A) Proposed mechanism for the reaction between the salicylaldehyde-containing probe (**1**) and Sph or Spa. During the reaction, the fluorescent dye “F” is covalently attached to the sphingolipid base, and separated from quencher “Q”, resulting in a highly fluorescent lipid product. (B) Chemical structure of probes synthesized in this study. (C) Fluorescence emission spectra of samples containing **1** (5 µM) after 24 h incubation in a DOPC liposome solution containing Sph, Spa, serine (Ser), sphingosine-1-phosphate (Sph1P), or buffer (control).

In designing our probe, we chose to use Bodipy FL as the fluorophore because its neutral charge and lipophilicity enhance cell permeability and partitioning into biological membranes.^23^ Additionally, Bodipy FL’s excitation (503 nm) and emission (509 nm) maxima are compatible with common microscope and fluorimeter filter sets, facilitating use. As a complementary quencher, we chose to use Black Hole Quencher-1 (BHQ-1) due to its high quenching efficiency at the 509 nm excitation maximum of Bodipy FL.^24^ In previous work, we determined that 4-(diethylamino)salicylaldehyde esters provided optimal hydrolytic stability under physiological conditions.^19^ Therefore, we designed the reactive core of our probe to include analogous 4-dialkylamino functionality. With these considerations in mind, we synthesized probe **1** (Figure 1B). Additionally, we synthesized a control probe **2**, which is identical to **1** except that it lacks the aldehyde necessary for selective reaction with terminal 1,2-amino alcohol groups and thus, should not react with sphingolipid bases in the cell (Figure 1B).^19,20^ Using vesicles as a model membrane, we initially tested the ability of compounds **1** and **2** to react with Sph and Spa under physiological conditions (pH 7.4, 37 °C). We found that **1** showed good stability (< 4% hydrolysis over 24 h) and reacted with both Sph and Spa to form the expected fluorescently labeled lipids (Figure S1-S2). Control probe **2** was completely stable under these conditions and, as expected, showed no reaction with Sph or Spa over 24 h (Figure S3). To determine if our probe is compatible with common cellular lipids, we incubated **1** in vesicles composed of several naturally abundant lipid species for 24 h (Figure S4).^25^ Under these conditions, we observed no reaction products between **1** and any of the tested lipids, although 1-palmitoyl-2-oleoyl-sn-glycero-3-phosphoethanolamine (POPE) did appear to slightly accelerate background hydrolysis of the probe. Having confirmed the reactivity of **1** toward Sph and Spa, we next quantified the fluorescence turn-on of **1** in the presence of these sphingolipid bases and potentially competing biomolecules. We found that when **1** (5 µM) was incubated with Sph (0.25 mM) or Spa (0.25 mM) at 37 °C for 24 h, a 3 and 2.5-fold fluorescent turn-on respectively was observed. However, when **1** was incubated with serine (2.50 mM) or sphingosine-1-phosphate (Sph1P) (0.25 mM) under the same conditions, there was no change in fluorescence as compared to untreated probe **1** (Figure 1C). These results agree with previous reports that, in the presence of phospholipid membranes, salicylaldehyde-modified lipids are only reactive toward molecules that are both lipophilic and contain 1,2-amino alcohol functionality.^19^ We were particularly encouraged to find that **1** did not react with Sph1P, which is identical in structure to sphingosine except that the primary alcohol is phosphorylated. The selective reaction and turn-on of our probe with Sph and Spa in membranes encouraged us to explore the utility of this tool in live cells.

As mentioned, Sph is roughly 10-fold more abundant than Spa in mammalian cells.^22^ Therefore, any observed response from probe **1** in live cells is likely attributable to Sph. For this reason, we chose to focus on Sph as the main analyte in our studies. To determine if **1** could react with Sph in live cells, we incubated HeLa cells with **1** (7.5 µM) for 2 h. We then exchanged cell media for new media containing varying concentrations of Sph. We found that after a 20 h incubation, cells treated with exogenous Sph showed a dose-dependent increase in fluorescence (Figure 2A-B). To ensure that the observed increase in fluorescence was due to the reaction of **1** with Sph, and not a biological effect of the added Sph, we performed the same experiment with control probe **2**. We found that when cells were incubated with **2** (7.5 µM) and then treated with Sph, no significant difference in fluorescence was observed between nontreated cells and those treated with Sph (Figure 2A and 2C). Furthermore, to confirm that the observed increase in cellular fluorescence was the result of the generation of a labeled sphingosine product and not the release of free Bodipy FL carboxylic acid (**BFL**) from **1**, we treated cells with the expected Bodipy FL-Sph product (**5**) and **BFL** (Figure S5A). We found that cells treated with **5** showed a membrane fluorescence pattern consistent with that observed in cells treated with **1** and sphingosine, while cells treated with **BFL** showed diffuse fluorescence, which did not stain cellular membranes (Figure S5B). These results indicate that **1** reacts with sphingosine in living cells to generate Bodipy FL-Sph in a dose-dependent manner.

**Figure 2.**
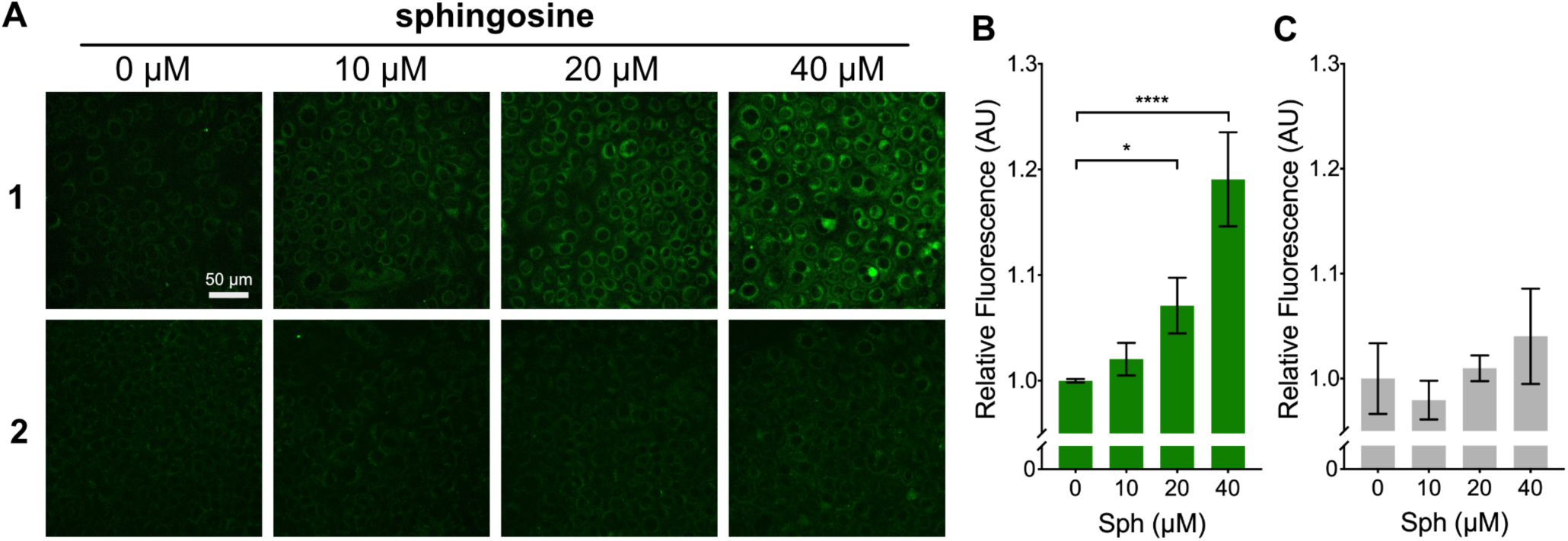
(A) Fluorescence microscopy images of HeLa cells treated with probe **1** and control probe **2** and exposed to a range of Sph concentrations (0-40 µM) for 20 h. (B and C) Quantified fluorescence response of **1** and **2** within large populations of cells after treatment with Sph (0-40 µM). Values are reported as means ± SD. Statistically significant changes in fluorescence are indicated as determined by one-way ANOVA: *P<0.05, ****P<0.0001.

Having demonstrated the compatibility of **1** in live cells through detection of exogenously added sphingosine, we next sought to apply this probe to the detection of endogenous levels of sphingosine in cells. We treated HeLa cells with **1** or **2** (20 µM) and imaged before and after a 16 h incubation. An increase in fluorescence (17.4%) was readily detected in cells treated with **1** over the course of the experiment (Figure 3A). In contrast, control probe **2** showed only a slight increase in fluorescence (4.7%), which was significantly less than probe **1** (Figure 3A-B). These results suggested that **1** is sensitive enough to detect native sphingosine in cells and may be used for probing differences in the sphingosine levels of cells.

**Figure 3.**
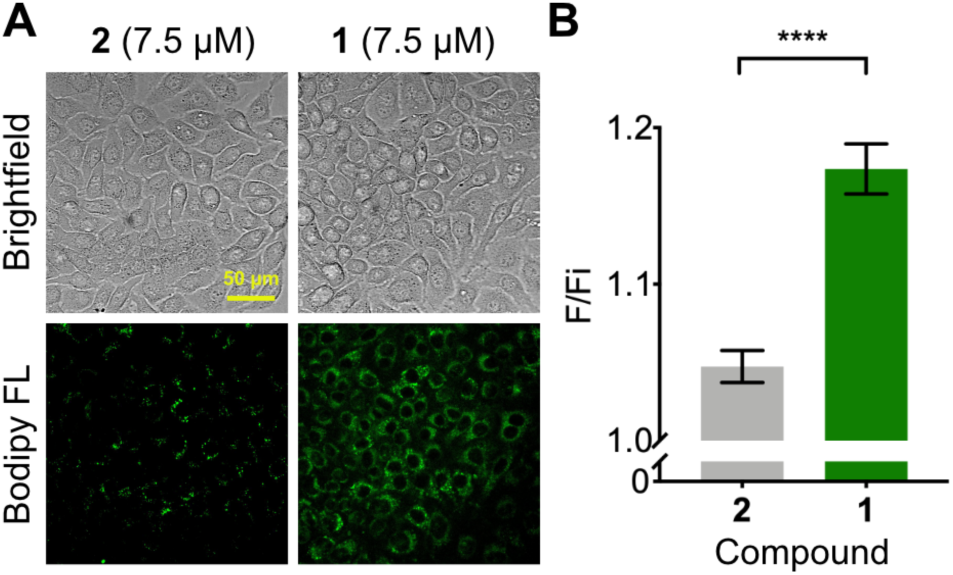
(A) Fluorescence microscopy images of HeLa cells treated with probe **1** and control probe **2** for 16 h. (B) Quantified fluorescence turn-on of **1** and **2** (final fluorescence [F]/ initial fluorescence [Fi]) within large populations of cells over 24 h. Values are reported as means ± SD. Significance was determined using an unpaired t-test: ****P<0.0001.

Changes in cellular sphingolipid levels occur in several diseases and often play a functional role in disease progression.^26–28^ Niemann-Pick Type C1 (NPC1) is a lysosomal storage disorder caused by a mutation in the *NPC1* gene, which encodes a large integral membrane protein (NPC1).^29^ The exact function of the NPC1 protein is not known, but when mutated in NPC1, it results in the accumulation of several lipids, including sphingosine, which plays a key role in promoting the disease phenotype.^10^ Patients with NPC1 feature a wide range of different neurological and systemic symptoms which differ from patient to patient making an accurate diagnosis difficult. Current diagnostic standards include filipin staining and DNA sequencing, but these tests are not always confirmative.^30^ Therefore, a probe for detecting increases in cellular sphingosine could be a valuable tool for the diagnosis of NPC1 and similar disorders. To test the ability of **1** to detect increased sphingosine in NPC1, we cultured fibroblasts derived from healthy and NPC1 patients. We incubated these cells in the presence of **1** (7.5 µM) for 24 h. We found that NPC1 cells showed a significantly higher fluorescence signal as compared to healthy fibroblasts (Figure 4). To determine if this difference in fluorescence correlated with the levels of Sph in the two cell lines, we quantified Sph in healthy fibroblasts and NPC1 cells by LCMS. In line with our fluorescence results, we found that the cell line derived from an NPC1 patient had higher levels of Sph compared to healthy fibroblasts (Figure S6). Analysis of the same samples showed that Spa levels were elevated in NPC1 cells, but, as expected, were roughly 10-fold less abundant than Sph, suggesting the majority of the fluorescence increase was due to increased Sph levels (Figure S6).

**Figure 4.**
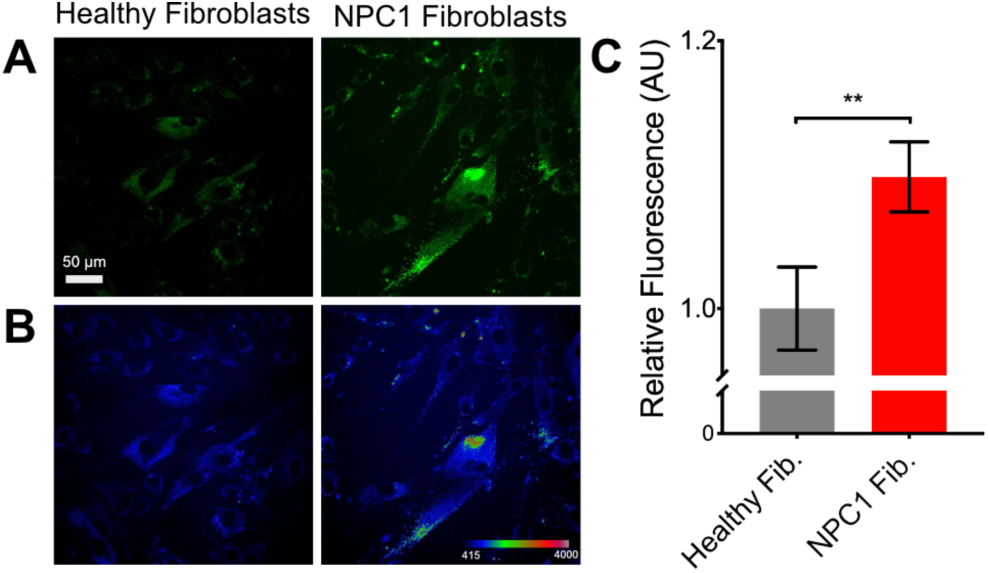
(A) Fluorescence microscopy images of Niemann-Pick disease type C1 (NPC1) patient-derived fibroblasts treated with probe **1** (7.5 µM) for 24 h. (B) Pixel intensity map of images in A. (C) Quantified fluorescence response of **1** within large populations of cells. Values are reported as means ± SD. Significance was determined using an unpaired t-test: **P<0.01.

In summary, we have developed the first chemical probe for the fluorogenic detection of Sph in living cells. The probe allows for the concentration-dependent detection of Sph in cultured mammalian cells using standard fluorescence microscopy techniques, which can be applied for the detection of increased Sph accumulation in NPC1 patient-derived cells. We envision that this straightforward approach for the detection of specific sphingolipids will facilitate their study in biology and may hold promise as a diagnostic tool for lipid storage disorders like NPC1 and Gaucher disease.^11^

## ASSOCIATED CONTENT

### Supporting Information

The Supporting Information is available free of charge on the ACS Publications website.

Supporting figures, detailed procedures, spectral data (PDF)

## AUTHOR INFORMATION

### Notes

The authors declare no competing financial interest.

## ACKNOWLEDGMENT

This study was supported by grants from the National Institute of Health (DP2DK111801-01S2 and R01 CA234245). Andrew K. Rudd thanks the NIH/NCI for his support though the Ruth L. Kirschstein National Research Service Award (T32 CA009523). Esther W. Lim thanks the NIH for her support (T32 EB009380).

---

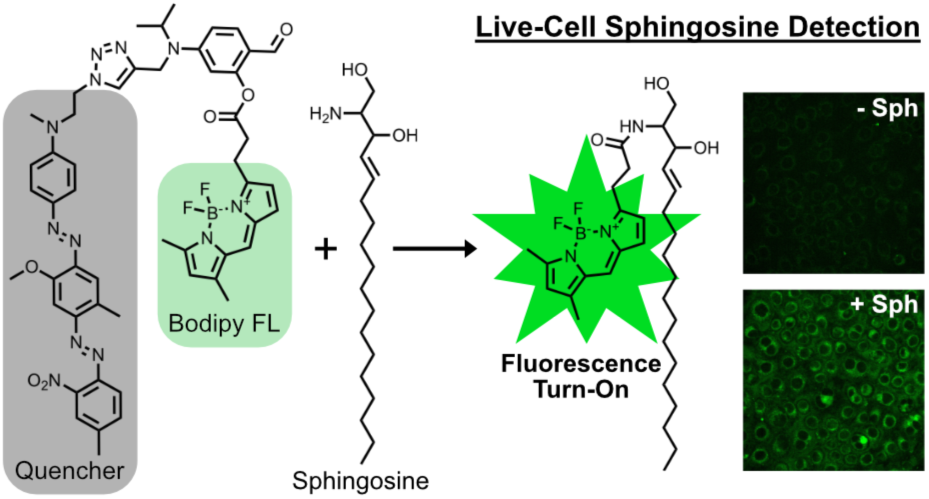

