## Supporting Information for "A Small Molecule Fluorogenic Probe for the Detection of Sphingosine in Living Cells"

#### Contents

General Procedures
Synthesis
Supporting Figures S1-S6
NMR Spectra
Mass Spectra
References

### General Procedures

**General procedure for synthesis of large unilamellar vesicles (LUVs)**
5 μmol of phospholipid in a 20 mg/mL $CHCl_3$ solution was added to a glass vial and dried into a film under a stream of nitrogen. 1 mL of buffer (1x PBS) was added to the vial and then the vial tumbled at RT for 1 hr. After tumbling, the formed solution was subjected to 10 freeze/thaw cycles using a dry ice/ethanol bath and a warm water bath with vortexing before every freeze step. The vesicle solution was then extruded 10 times through a 100 nm polycarbonate filter (Whatman, GE Healthcare Bio-Sciences) using an Avanti Mini-Extruder (Avanti Polar Lipids) to produce LUVs.

**Fluorescence measurements in DOPC vesicles**
Solutions of 5 mM 1,2-dioleoyl-sn-glycero-3-phosphocholine (DOPC) vesicles with or without sphingosine (250 μM), sphinganine (250 μM), sphingosine-1-phosphate (250 μM), and serine (2.5 mM) were treated with Compound **1** (200 μM stock in PBS) at a final concentration of 5 μM. Reactions were incubated on a tube rotator at 37 ºC for 24 h. Fluorescence emission spectra from 500-560 nm (ex. 480 nm) for each sample were obtained on a Jasco FP-8500 spectrofluorometer.

**Probe reaction in DOPC vesicles**
Solutions of 5 mM DOPC vesicles in PBS with or without sphingosine (1 mM) or sphinganine (1 mM) were treated with Compound **1** (100 μM). Reactions were incubated on a tube rotator at 37 ºC for 24 h. Samples were analyzed using HPLC Method A before and after 24 h incubation.

**Probe stability and reactivity in naturally abundant lipid vesicles**
Solutions of 5 mM 1-palmitoyl-2-oleoyl-sn-glycero-3-phosphocholine (POPC) vesicles in PBS with or without 1-palmitoyl-2-oleoyl-sn-glycero-3-phosphoethanolamine (POPE) (1 mM), 1-palmitoyl-2-oleoyl-sn-glycero-3-phospho-L-serine (POPS) (1 mM), or cholesterol (Chol) (1 mM) were treated with Compound **1** (100 μM). Reactions were incubated on a tube rotator at 37 ºC for 24 h. Samples were analyzed using HPLC Method A before and after 24 h incubation.

**Cell culture**
Healthy Human Fibroblasts (GM05399) and Infantile Neuronal Ceroid Lipofuscinosis Human Fibroblasts (GM20389) were obtained from Coriell Institute (Camden, NJ). HeLa S3 (CCL-2.2) cells were obtained from ATCC (Manassas, VA). Cells were maintained in DMEM supplemented with Penicillin (50 units/mL), Streptomycin (50 μg/mL) and 10 % FBS (Healthy Fibroblasts and HeLas) or 15 % FBS (INCL Fibroblasts) at 37 ºC, 5 % $CO_2$.

**Probe dose-response in HeLa cells**
HeLa cells were plated in an 8-well Lab-Tek chamber slide (Sigma-Aldrich) at a density of 40,000 cells/well and allowed to attach overnight. Media was removed and then 500 μL of media containing 7.5 μM **1** or **2** was added to each well and cells incubated at 37 ºC, 5 % $CO_2$ for 2 h. Media in each well was then exchanged for 500 μL of media containing the indicated concentrations of sphingosine or sphinganine and the cells incubated at 37 ºC, 5 % $CO_2$ for 20 h before imaging.

**Imaging of Bodipy FL carboxylic acid and sphingolipid products in HeLa cells**

HeLa cells were plated in an 8-well Lab-Tek chamber slide (Sigma-Aldrich) at a density of 40,000 cells/well and allowed to attach overnight. Media was removed and then 500 µL of media containing 5 µM Bodipy FL carboxylic acid, **5,** or **6** was added to each well and cells incubated at 37 ºC, 5 % $CO_2$ for 1 h before imaging.

**Detection of endogenous sphingosine/sphinganine in HeLa cells**

HeLa cells were plated in an 8-well Lab-Tek chamber slide (Sigma-Aldrich) at a density of 40,000 cells/well and allowed to attach overnight. Media was removed and each well washed once with OptiMEM media (- FBS) before adding 500 µL OptiMEM (- FBS) containg 20 µM **1** or **2**. Cells were incubated at 37 ºC, 5 % $CO_2$ for 4 h before exchanging media in each well for growth media. Cells were then maintained at 37 ºC, 5 % $CO_2$ for 16 h while imaging.

**Detection of sphingosine/sphinganine in Healthy and NPC1 fibroblasts**

Healthy and NPC1 fibroblast cells were plated in an 8-well Lab-Tek chamber slide (Sigma-Aldrich) at a density of 20,000 cells/well and allowed to attach overnight. Media was removed and 500 µL DMEM (15 % FBS) containing 7.5 µM **1** was added to each well. Cells were incubated at 37 ºC, 5 % $CO_2$ for 24 h before imaging.

**Quantification of cellular fluorescence**

Using ImageJ,[1] the mean fluorescence of individual cells was measured within 4 images taken across at least 2 biological replicates for each condition. The mean cellular fluorescence intensity was then calculated for each of the 4 images and data reported as the mean of these 4 means ± SD.

**Sphingosine/sphinganine extraction and measurement in Healthy and NPC1 fibroblasts**

Cells seeded in 6-well plates were extracted with 250 µL -20 °C methanol, 150 µL of saline, 100 µL of water and spiked with a mixture of deuterated standards (sphingosine-d7, sphinganine-d7 (Avanti Lipids)) of a known concentration as internal control. 500 µL of chloroform was added and the samples were vortexed for 5 min followed by centrifugation at 4 °C for 5 min at 16 000 x *g*. The organic phase was collected and 1 µL of formic acid was added to the remaining polar phase which was re-extracted with 500 µL of chloroform. Combined organic phases were dried and the pellet was resuspended in 100 µL of buffer containing 100% methanol, 1 mM ammonium formate and 0.2% (v/v) formic acid. Quantification of these species were determined via liquid chromatography mass-spectrometry (Agilent 6460 QQQ). The sphingoid bases were separated on C8 column (Spectra 3 µm C8SR 150 x 3mm ID, Peeke Scientific, CA) as previously described.[2] Mobile phase A was composed of 100% HPLC grade water containing 2 mM ammonium formate and 0.2% (v/v) formic acid and mobile phase B consisted of 100% methanol containing 0.2% (v/v) formic acid and 1 mM ammonium formate. The gradient elution program consisted of holding at 82% B for 3 min, up to 90% B over 1 min and linearly increasing to 99% B over 14 min maintaining it for 7 min and down to 82% B over 2 min. The 82% mobile phase B was maintained for 3 min followed by a re-equilibration for 10 min. The capillary voltage was set to 3.5 kV, the drying gas temperature was 350 °C, the drying gas flow rate was 10 L/min, and the nebulizer pressure was 60 psi. Sphingoid base species were analyzed by SRM of the transition from precursor to product ions at associated optimized collision energies and fragmentor voltages (Table below). Abundances of these species were then quantified from spiked internal standards.

| Sphingolipid species | Parent | Daughter | Fragmentor | CE |
|---|---|---|---|---|
| SA d18:0 | 302.4 | 284.4 | 120 | 9 |
| SA-d7 d18:0 | 309.4 | 291.5 | 120 | 9 |
| SO d18:1 | 300.3 | 282.4 | 112 | 9 |
| SO-d7 d18:0 | 307.3 | 289.4 | 112 | 9 |

**Microscopy**
Imaging was performed on an Axio Observer Z1 inverted microscope (Carl Zeiss Microscopy Gmb, Germany) with Yokogawa CSU-X1 spinning disk confocal unit using a 20x, 0.8 NA objective to an ORCA-Flash4.0 V2 Digital CMOS camera (Hamamatsu, Japan). Fluorophores were excited with a diode laser (488 nm; 30 mW). Where indicated, cells were maintained at 37 ºC, 5% $CO_2$ using an Okolab stagetop incubator (Italy). Images were acquired using Zen Blue software (Carl Zeiss) and processed using Image J.

**HPLC**
*Method A*
Samples were analyzed or purified on an Agilent 1100 series HPLC equipped with a diode array detector (DAD) using a Zorbax SB-C18 column (Agilent) with HPLC grade $H_2O$ and MeOH each containing 0.1% formic acid. Samples were eluted using a gradient: 75:25 to 95:5 MeOH:$H_2O$ from 0-5 min, 95:5 MeOH:$H_2O$ from 5-17 min.

**Statistical Methods**
All statistical analysis was performed using Prism 7 software. Where shown, error bars represent standard deviation.

### Synthesis

***Synthesis of Sph/Spa Fluorescent Probe 1***

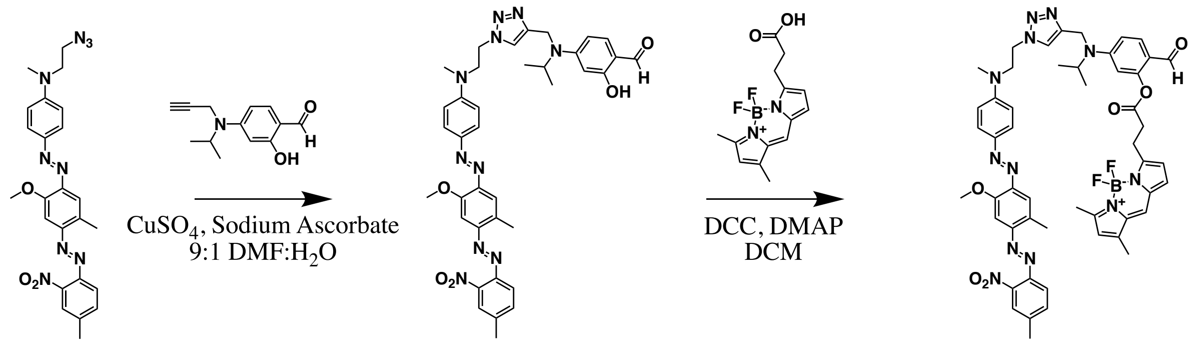


**BHQ-1-$N_3$** **3** **1**

**2-hydroxy-4-(isopropyl(BHQ-1-$N_3$)amino)benzaldehyde (3).** BHQ-1-$N_3$[3] (11.3.mg, 0.023 mmol) and 2-hydroxy-4-(isopropyl(prop-2-yn-1-yl)amino)benzaldehyde[4] (7.6 mg, 0.035 mmol) were dissolved in 1.5 mL 9:1 DMF:$H_2O$. $CuSO_4$ pentahydrate (0.6 mg, 0.002 mmol) and sodium ascorbate (0.9 mg, 0.005 mmol) were added and the reaction stirred at 50 ºC under inert atmosphere for 12 h. Solvent was removed by rotary evaporation, the residue dissolved in $CH_3CN$ and purified using HPLC method A. Solvent was removed by rotary evaporation to yield **3** as a dark red solid (11.6 mg, 71 % yield). $^1$H NMR (500 MHz, Chloroform-d) δ 11.54 (s, 1H), 9.54 (s, 1H), 7.86 (d, J = 8.8 Hz, 2H), 7.72 (s, 1H), 7.67 (d, J = 8.2 Hz, 1H), 7.58 (s, 1H), 7.49 (d, J = 8.1 Hz, 1H), 7.41 (s, 1H), 7.24 (d, J = 8.9 Hz, 1H), 7.11 (s, 1H), 6.60 (d, J = 8.8 Hz, 2H), 6.29 (d, J = 8.8 Hz, 1H), 6.15 (s, 1H), 4.62 – 4.53 (m, 4H), 4.29 – 4.19 (m, 1H), 4.03 (s, 3H), 3.96 (t, 2H), 2.83 (s, 3H), 2.72 (s, 3H), 2.53 (s, 3H), 1.19 (d, J = 6.5 Hz, 6H). $^{13}$C NMR (125 MHz, Chloroform-d) δ 192.5, 164.1, 154.9, 154.9, 151.3, 150.5, 147.5, 146.6, 145.2, 145.0, 143.4, 141.7, 135.4, 133.5, 132.9, 125.7, 124.3, 122.9, 119.0, 119.0, 112.2, 111.4, 105.3, 99.4, 98.3, 56.3, 52.6, 48.8, 47.7, 40.3, 39.0, 21.3, 20.2, 16.7. MS (ESI) calculated $[M+H]^+$: 705.3, found: 705.3.

**2-(Bodipy FL)-4-(isopropyl(BHQ-1-$N_3$)amino)benzaldehyde (1).** BDP FL carboxylic acid [Lumiprobe Hunt Valley, MD] (1.6 mg, 5.5 µmol) was dissolved in 200 µL dry DCM. 1 M N,N'-Dicyclohexylcarbodiimide in DCM (5.5 µL, 5.5 µmol) was added and the reaction stirred at rt for 10 min. **3** (3.0 mg, 4.3 µmol) and 4-(Dimethylamino)pyridine (0.8 mg, 6.4 µmol) were dissolved in 300 µL dry DCM and added to the reaction before stirring under inert atmosphere at rt for 4 h. The reaction was filtered through a 0.2 µm filter and dried by rotary evaporation. The residue was dissolved in $CH_3CN$ and purified using HPLC method A. Solvent was removed by rotary evaporation to yield **1** as a dark red solid (2.8 mg, 67 % yield). MS (ESI) calculated $[M+H]^+$: 979.4, found: 979.4. $^1$H NMR (500 MHz, dimethyl sulfoxide-d6) δ 9.65 (s, 1H), 8.35 (s, 1H), 7.94 (m, 2H), 7.75 (d, J = 8.1 Hz, 1H), 7.70 (m, 2H), 7.60 (d, J = 9.0, 1H), 7.50 (s, 1H), 7.28 (s, 1H), 7.09 (d, J = 4.0, 1H), 6.73 (m, 3H), 6.55 (s, 1H), 6.47 (d, J = 4.0, 1H), 6.31 (s, 1H), 4.57 (t, J = 5.8 Hz, 2H), 4.53 (s, 2H), 4.26 (m, 1H), 3.91 (m, 5H), 3.22 (t, J = 8.0, 2H), 3.06 (t, J = 8.0, 2H), 2.77 (s, 3H), 2.62 (s, 3H), 2.53* (s, 3H), 2.46 (s, 3H), 2.25 (s, 3H), 1.13 (d, J = 6.5, 6H). *peak obscured by residual dimethyl sulfoxide solvent peak but predicted based on equivalent $CH_3$ peak in precursor (**3**). $^{13}C$ NMR (126 MHz, dimethyl sulfoxide-d6) δ 186.9, 170.8, 159.9, 156.0, 154.5, 153.8, 153.1, 151.4, 150.3, 146.4, 145.3, 144.6, 143.9, 142.7, 142.3, 134.8, 134.0, 133.2, 133.1, 132.3, 128.8, 128.8, 125.6, 125.6, 125.3, 124.4, 123.6, 120.6, 120.2, 118.6, 116.6, 111.5, 109.8, 105.7, 105.7, 99.1, 79.2, 55.9, 51.6, 48.4, 47.1, 38.1, 32.1, 23.3, 20.8, 19.8, 16.3, 14.6, 11.1.

***Synthesis of Control Probe 2***

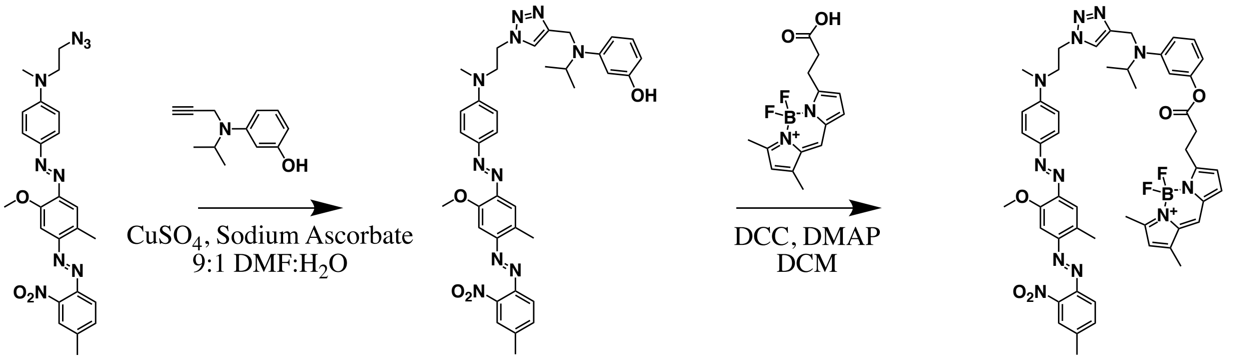


**BHQ-1-N$_3$** **4** **2**

**3-(isopropyl(BHQ-1-N$_3$)amino)phenol (4).** BHQ-1-N$_3$[3] (9.3 mg, 0.019 mmol) and 3-(isopropyl(prop-2-yn-1-yl)amino)phenol[4] (5.4 mg, 0.029 mmol) were dissolved in 1.5 mL 9:1 DMF:$H_2O$. $CuSO_4$ pentahydrate (0.5 mg, 0.002 mmol) and sodium ascorbate (0.8 mg, 0.004 mmol) were added and the reaction stirred at 50 ºC under inert atmosphere for 12 h. Sovlvent was removed by rotary evaporation, the residue dissolved in $CH_3CN$ and purified using HPLC method A. Solvent was removed by rotary evaporation to yield **3** as a dark red solid (8.7 mg, 67 % yield). 1H NMR (500 MHz, Chloroform-d) δ 7.86 (d, J = 8.9 Hz, 2H), 7.73 (s, 1H), 7.67 (d, J = 8.2 Hz, 1H), 7.59 (s, 1H), 7.49 (d, J = 8.2 Hz, 1H), 7.42 (s, 1H), 7.08 (s, 1H), 7.04 – 6.97 (m, 1H), 6.53 (d, J = 9.0 Hz, 2H), 6.30 (d, J = 8.6 Hz, 1H), 6.20 (d, J = 6.0 Hz, 2H), 4.54 (t, J = 5.5 Hz, 2H), 4.46 (s, 2H), 4.10 – 4.05 (m, 1H), 4.05 (s, 3H), 3.93 (t, J = 5.6 Hz, 2H), 2.74 (s, 3H), 2.72 (s, 3H), 2.53 (s, 3H), 1.13 (d, J = 6.6 Hz, 6H). 13C NMR (125 MHz, Chloroform-d) δ 157.1, 154.7, 151.2, 150.6, 149.8, 147.5, 145.0, 144.9, 143.4, 141.7, 133.6, 133.5, 133.1, 130.0, 125.9, 124.4, 123.2, 119.0, 119.0, 111.5, 106.0, 104.3, 101.0, 99.3, 56.2, 52.6, 48.9, 47.5, 40.0, 38.9, 21.4, 20.1, 16.8. MS (ESI) calculated $[M+H]^+$: 677.3, found: 677.3.

**3-(isopropyl(BHQ-1-N$_3$)amino)phenyl Bodipy FL (2).** BDP FL carboxylic acid [Lumiprobe, Hunt Valley, MD] (1.9 mg, 6.4 µmol) was dissolved in 200 µL dry DCM. 1 M N,N'-Dicyclohexylcarbodiimide in DCM (6.4 µL, 6.4 µmol) was added and the reaction stirred at rt for 10 min. **4** (4.3 mg, 6.4 µmol) and 4-(Dimethylamino)pyridine (0.8 mg, 6.4 µmol) were dissolved in 300 µL dry DCM and added to the reaction before stirring under inert atmosphere at rt for 12 h. The reaction was filtered through a 0.2 µm filter and dried by rotary evaporation. The residue was dissolved in 1:1 MeOH:$CH_3CN$ and purified using HPLC method A. Solvent was removed by rotary evaporation to yield **1** as a dark red solid (0.6 mg, 10 % yield). MS (ESI) calculated $[M+H]^+$: 951.4, found: 951.4. $^1H$ NMR (500 MHz, dimethyl sulfoxide-d6) δ 7.95 (s, 1H), 7.87 (s, 1H), 7.71 (m, 4H), 7.50 (s, 1H), 7.28 (s, 1H), 7.12 (d, J = 8.2 Hz, 1H), 7.09 (m, 1H), 6.73 (d, J = 9.3 Hz, 2H), 6.56 (d, J = 8.6 Hz, 1H), 6.42 (m, 2H), 6.33 (d, J = 7.9 Hz, 1H), 6.31 (s, 1H), 4.55 (t, J = 5.9 Hz,

2H), 4.36 (s, 2H), 4.07 (m, 1H), 3.91 (m, 5H), 3.19 (t, J = 7.4 Hz, 2H), 2.97 (t, J = 7.4 Hz, 2H), 2.75 (s, 3H), 2.62 (s, 3H), 2.53* (s, 3H), 2.46 (s, 3H), 2.25 (s, 3H), 1.11 (d, J = 6.4 Hz, 6H). *peak obscured by residual dimethyl sulfoxide solvent peak but predicted based on equivalent $CH_3$ peak in precursor (**4**). $^{13}C$ NMR (126 MHz, Chloroform-d) δ 171.4, 160.8, 156.5, 155.0, 152.0, 151.3, 150.8, 149.6, 148.0, 147.6, 145.2, 144.2, 143.6, 141.8, 135.4, 133.7, 133.4, 133.1, 130.0, 129.2, 128.4, 128.1, 126.0, 125.4, 124.9, 124.5, 124.1, 123.3, 120.7, 119.2, 119.1, 116.9, 111.5, 110.9, 109.8, 106.9, 99.4, 56.4, 52.7, 51.1, 48.9, 47.6, 40.4, 39.0, 33.7, 31.1, 29.9, 24.1, 21.6, 21.5, 20.1, 16.9, 15.1, 11.5.

***Synthesis of Bodipy FL-Sph 5 and Bodipy FL-Spa 6***

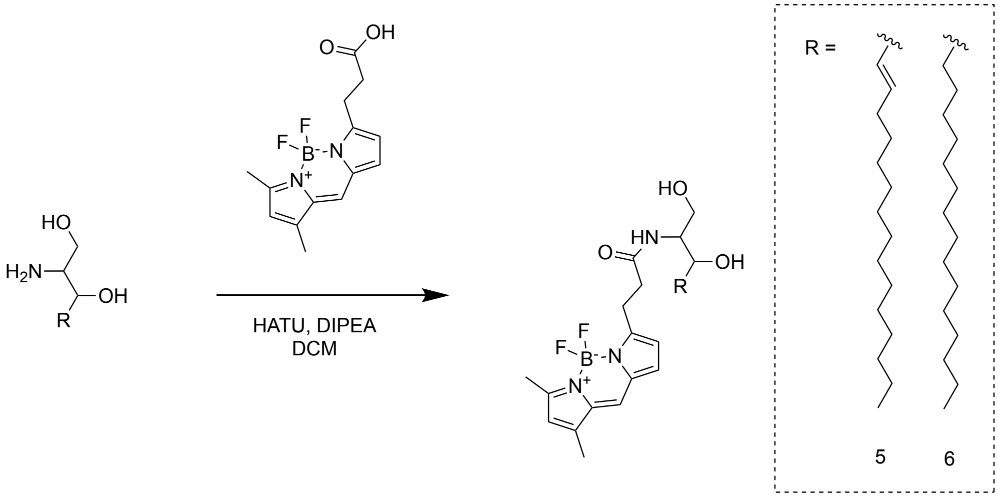


**Bodipy FL-sphingosine (5).** BDP FL carboxylic acid [Lumiprobe, Hunt Valley, MD] (3.5 mg, 12.0 µmol) was dissolved in 200 µL dry DCM and stirred at 0 ºC for 10 min. HATU (5.0 mg, 13.2 µmol) and DIPEA (8.4 µL, 47.9 µmol) were then added and the reaction stirred at 0 ºC for 10 min. D-erythro-sphingosine (3.6 mg, 12.0 mmol) [Avanti Polar Lipids, Alabaster, AL] was then added and the reaction stirred at rt for 1 h. Solvent was removed by rotary evaporation and the residue dissolved in MeOH and purified using HPLC method A. Solvent was removed by rotary evaporation to yield **5** as a red solid (5.0 mg, 73 % yield). MS (ESI) calculated $[M+Na]^+$: 596.4, found: 596.5. $^1H$ NMR (500 MHz, Chloroform-d) δ 7.09 (s, 1H), 6.89 (d, J = 3.9 Hz, 1H), 6.63 (bs, 1H), 6.31 (d, J = 3.9 Hz, 1H), 6.13 (s, 1H), 5.73 (m, 1H), 5.45 (m, 1H), 4.25 (m, 1H), 3.84 (m, 2H), 3.63 (m, 1H), 3.30 (t, J = 7.4 Hz, 2H), 2.75 (t, J = 7.4 Hz, 2H), 2.59 (s, 3H), 2.25 (s, 3H), 2.00 (q, J = 7.2 Hz, 2H), 1.38-1.18 (m, 22H), 0.87 (t, J = 7.1 Hz, 3H). $^{13}C$ NMR (125 MHz, Chloroform-d) δ 173.2, 161.1, 156.5, 144.5, 135.5, 134.4, 133.4, 128.5, 128.1, 124.0, 120.9, 117.5, 74.3, 62.3, 55.1, 35.8, 32.4, 32.1, 29.9, 29.8, 29.6, 29.5, 29.4, 29.2, 25.1, 22.8, 15.2, 14.3, 11.5

**Bodipy FL-sphinganine (6).** BDP FL carboxylic acid [Lumiprobe, Hunt Valley, MD] (6.1 mg, 20.9 µmol) was dissolved in 200 µL dry DCM and stirred at 0 ºC for 10 min. HATU (8.7 mg, 23.0 µmol) and DIPEA (14.6 µL, 83.5 µmol) were then added and the reaction stirred at 0 ºC for 10 min. D-erythro-sphinganine (6.3 mg, 20.9 µmol) [Avanti Polar Lipids, Alabaster, AL] was then added and the reaction stirred at rt for 1 h. Solvent was removed by rotary evaporation and the residue dissolved in MeOH and purified using HPLC method A. Solvent was removed by rotary evaporation to yield **6** as a red solid (9.7 mg, 81 % yield). MS (ESI) calculated $[M+Na]^+$: 598.4, found: 598.5. $^{1}H$ NMR (500 MHz, Chloroform-d) δ 7.09 (s, 1H), 6.89 (d, J = 3.9 Hz, 1H), 6.80 (bs, 1H), 6.32 (d, J = 3.9 Hz, 1H), 6.13 (s, 1H), 3.90 (m, 1H), 3.76 (bs, 1H), 3.68 (m, 2H), 3.29 (t, J = 7.0 Hz, 2H), 2.75 (t, J = 7.0 Hz, 2H), 2.55 (s, 3H), 2.25 (s, 3H), 1.43 (m, 2H), 1.24 (m, 26H), 0.87 (t, 7.1 Hz, 3H). $^{13}C$ NMR (125 MHz, Chloroform-d) δ 172.9, 161.1, 156.5, 144.5, 135.5, 133.4, 128.1, 124.0, 120.9, 117.5, 73.8, 62.3, 54.6, 35.7, 34.3, 32.1, 29.9, 29.8, 29.5, 26.1, 25.1, 22.8, 15.2, 14.3, 11.5.

### Supporting Figures

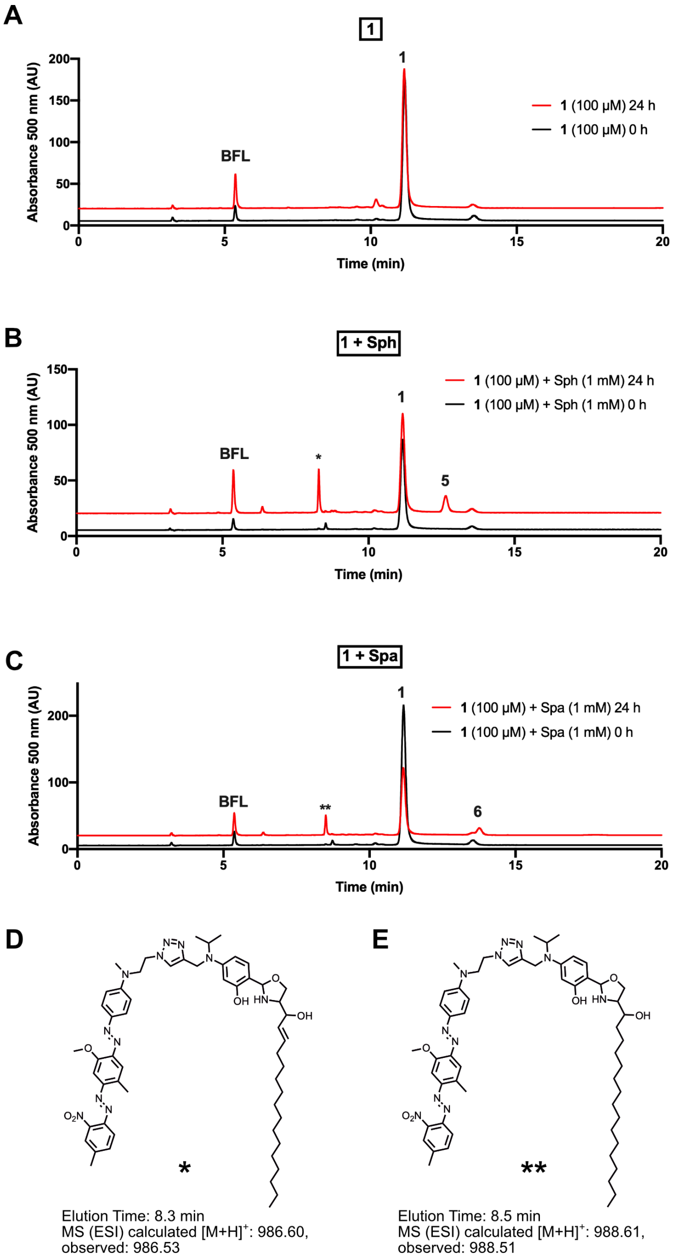


**Figure S1.** Probe **1** reacts with Sph and Spa to form Bodipy FL-labeled sphingolipid products **5** and **6**. (**A-C**) HPLC A500 traces before and after incubation of **1** (100 µM) alone (A), with Sph (1 mM) (B), or with Spa (1 mM) (C). Under reaction conditions, small amounts of Bodipy FL carboxylic acid **BFL** are generated due to background hydrolysis. (**D-E**) Side products were identified upon reaction of **1** with Sph or Spa. These were identified as adducts between the 2-hydroxy-4-(isopropyl(BHQ-1-$N_3$)amino)benzaldehyde **3** (which is released during the formation of the fluorescent lipid product) and Sph (D) or Spa (E). Analogous side products were previously reported for the reaction between Sph and other salicylaldehyde esters.[5] Peaks were identified from known standards of **1**, **5**, **6**, **BFL** (Bodipy FL carboxylic acid) or by MS analysis of purified peaks * (D) and ** (E) (see *Mass Spectra*).

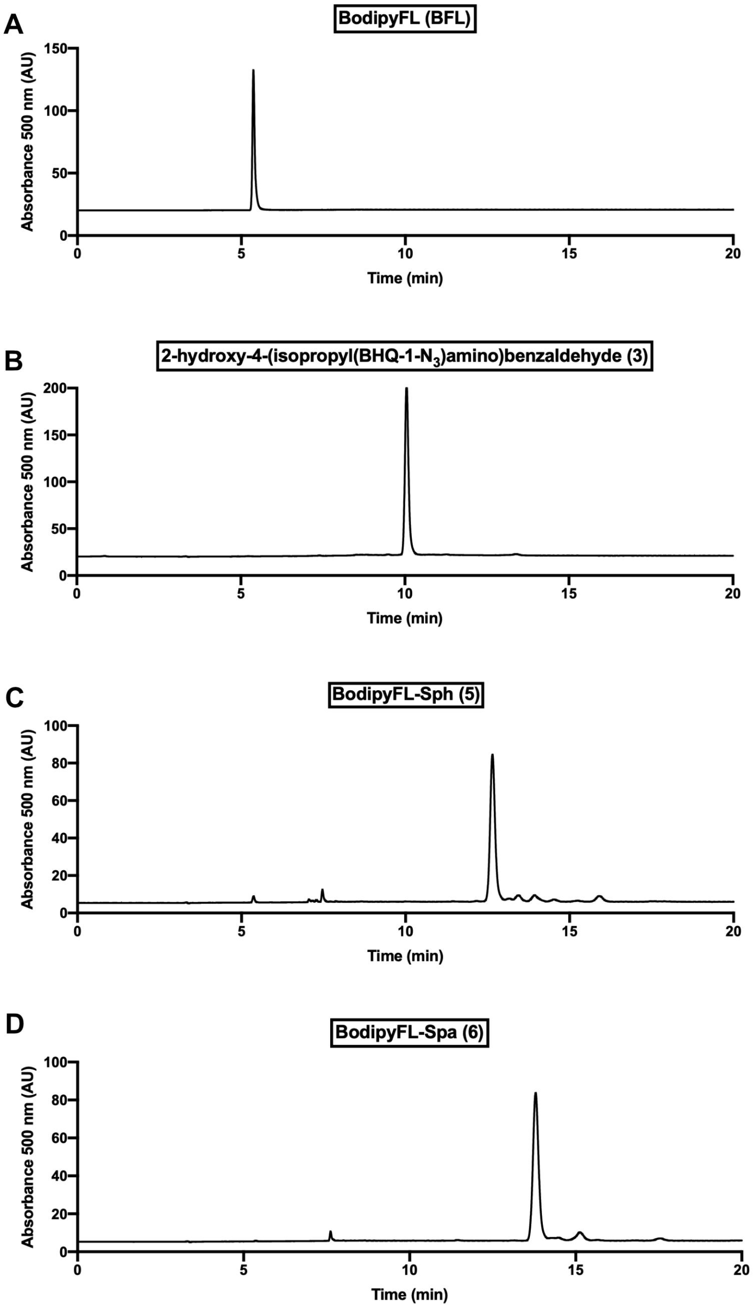


**Figure S2.** HPLC A500 traces of known standards of Bodipy FL carboxylic acid (**A**), 2-hydroxy-4-(isopropyl(BHQ-1-$N_3$)amino)benzaldehyde **3** (**B**), Bodipy FL-Sph **5** (C), and Bodipy FL-Spa **6** (D).

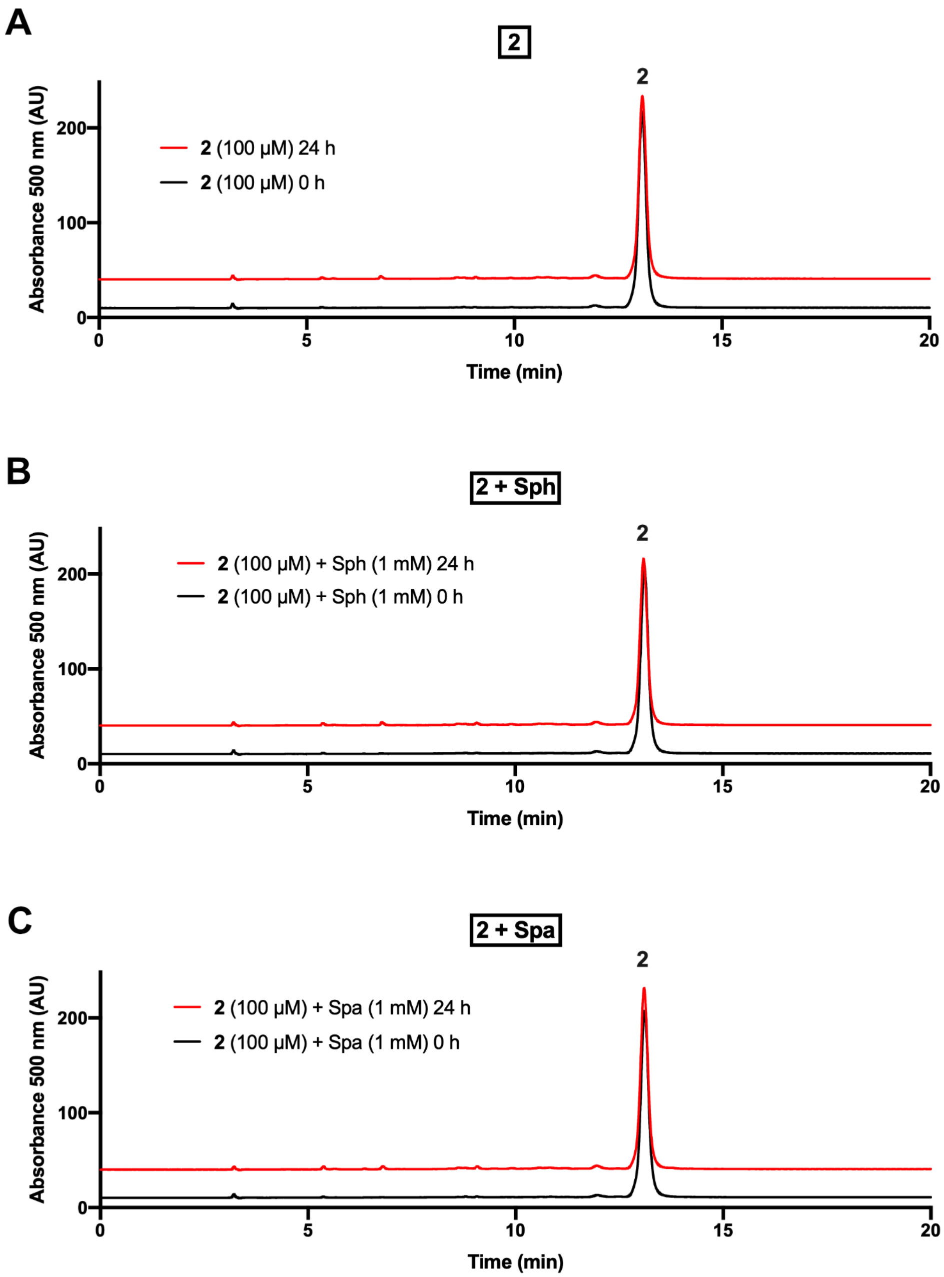


**Figure S3.** Probe **2** is unreactive toward Sph and Spa. (**A-C**) HPLC A500 traces before and after incubation of **2** (100 µM) alone (A), with Sph (1 mM) (B), or with Spa (1 mM) (C).

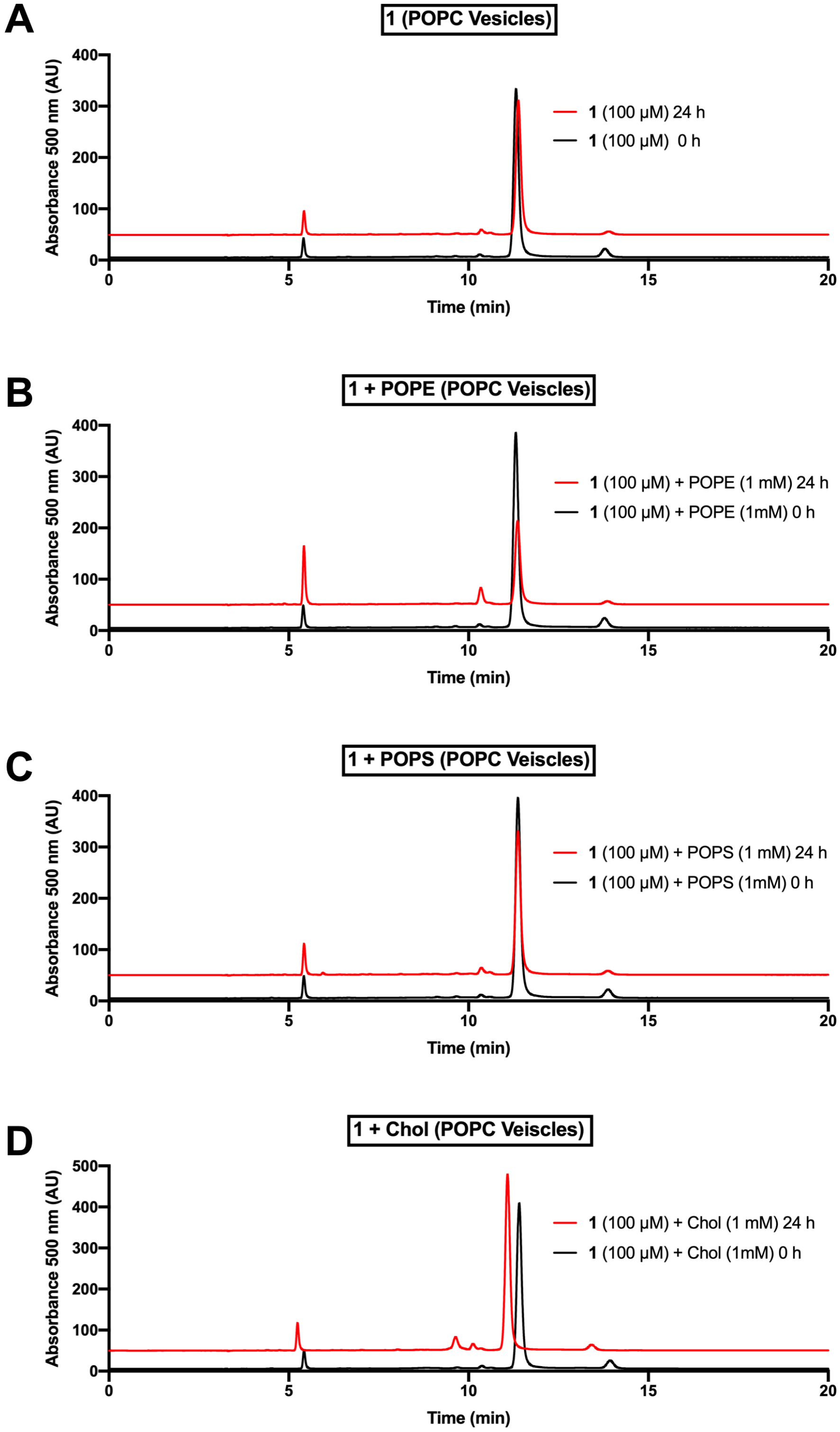


**Figure S4.** Probe **1** stability and reactivity in the presence of naturally abundant lipid species. (**A-D**) HPLC A500 traces before and after incubation of **1** (100 µM) in POPC Vesicles (5 mM) Alone (A), with POPE (1 mM) (B), POPS (1 mM) (C), or Cholesterol (D).

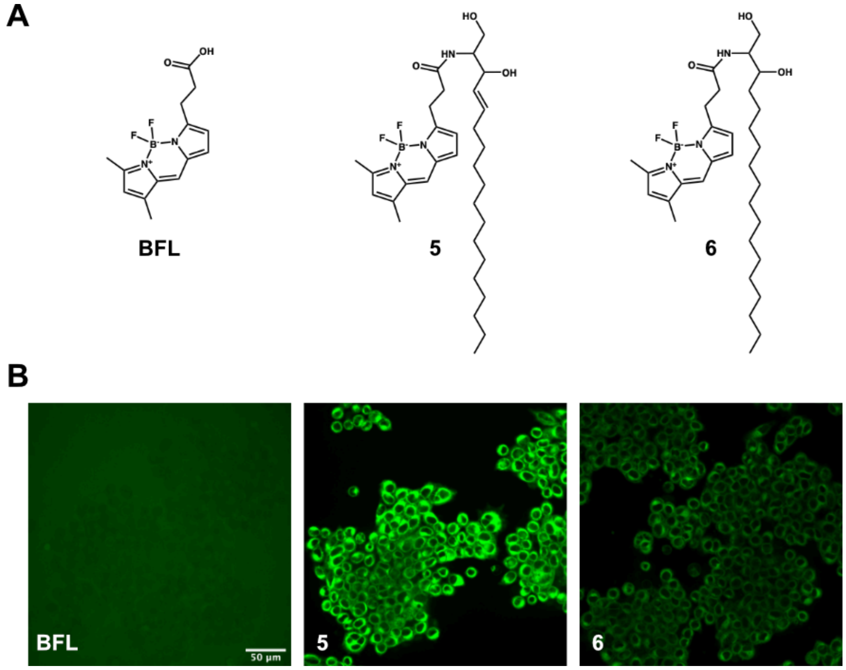


**Figure S5.** (**A**) Chemical structures of Bodipy FL carboxylic acid (**BFL**), Bodipy FL-sphingosine (**5**), and Bodipy FL-sphinganine (**6**). (**B**) Fluorescence microscopy images of HeLa cells after a 1 h incubation with 5 µM **BFL**, **5**, and **6**. **BFL** does not partition into biological membranes while both Bodipy FL-labeled sphingoid bases **5** and **6** show membrane localization similar to that observed in cells treated with compound **1**.

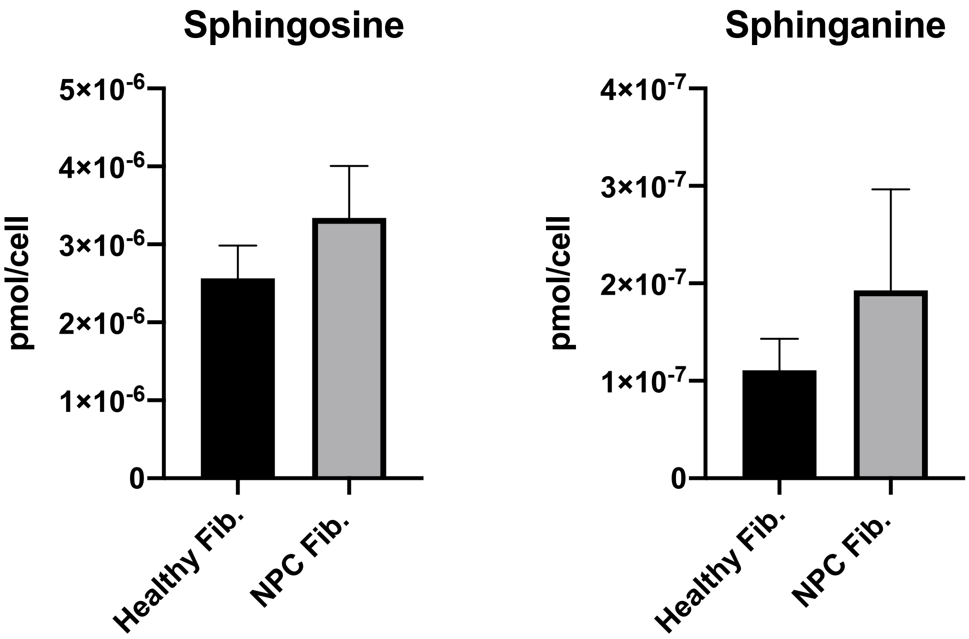


**Figure S6.** LCMS detection of Sph and Spa in Healthy and NPC1 fibroblasts. Values are reported as means ± SD.

### NMR Spectra

#### Compound 1 $^1$H and $^{13}$C NMR

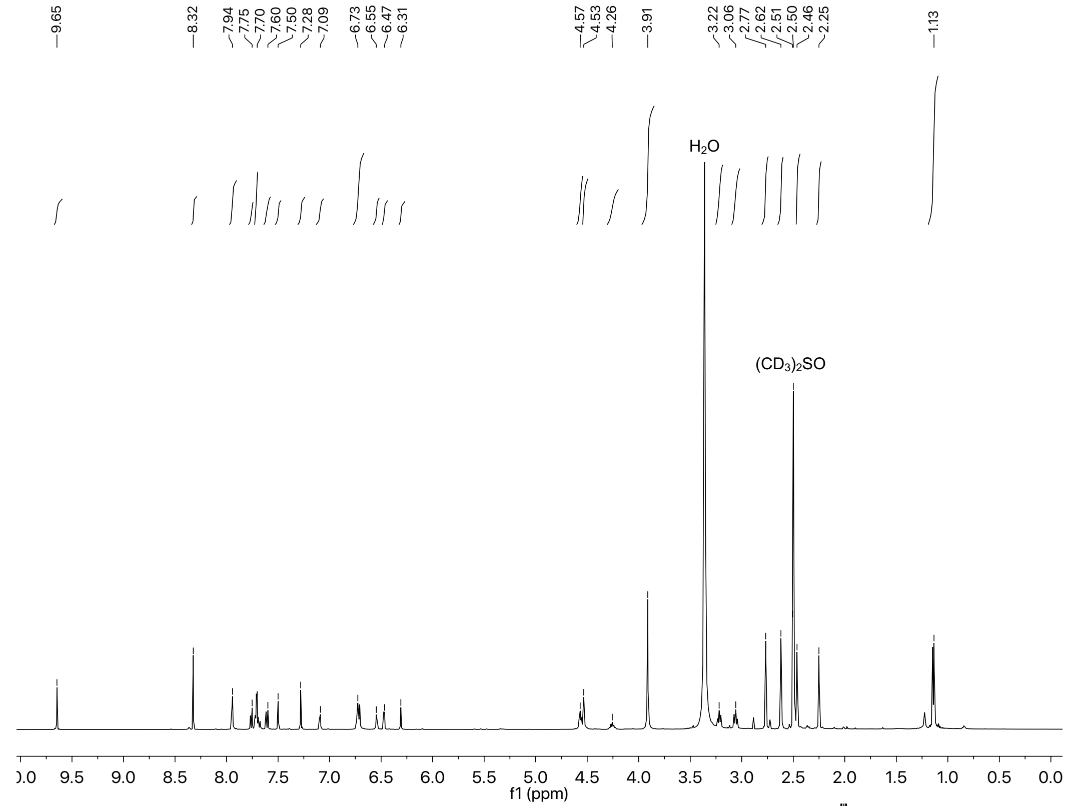

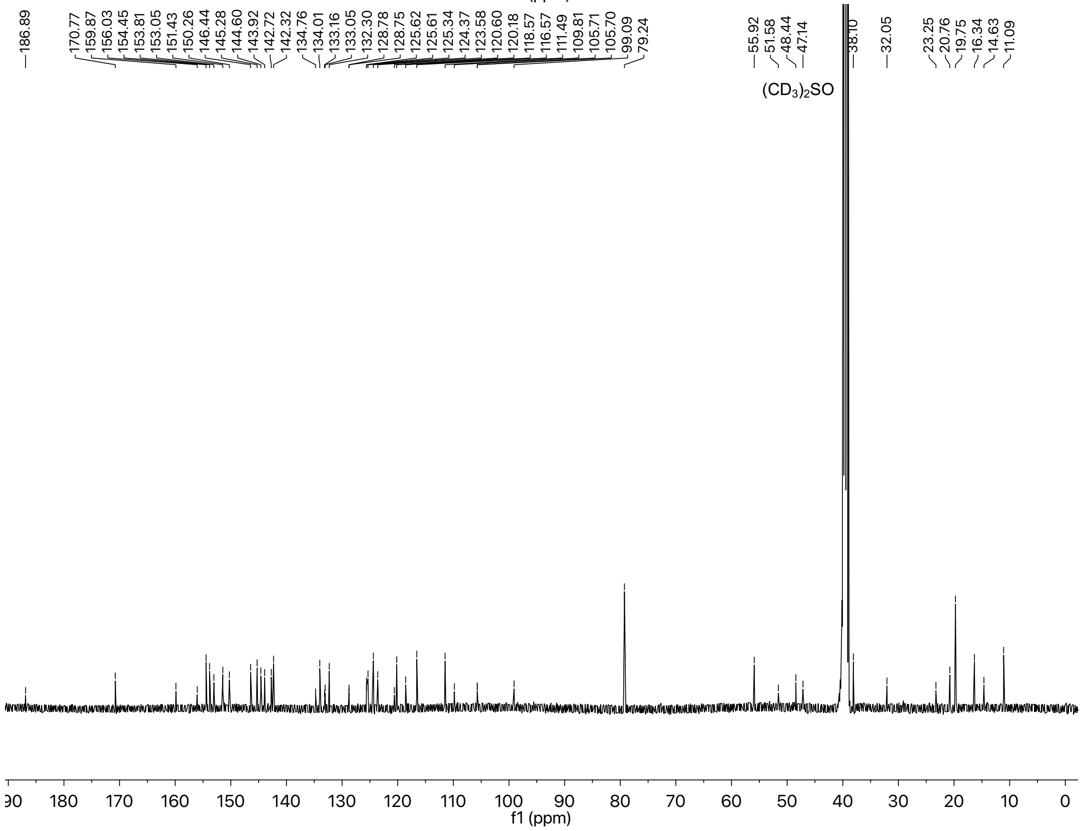

#### Compound 2 $^{1}H$ and $^{13}C$ NMR

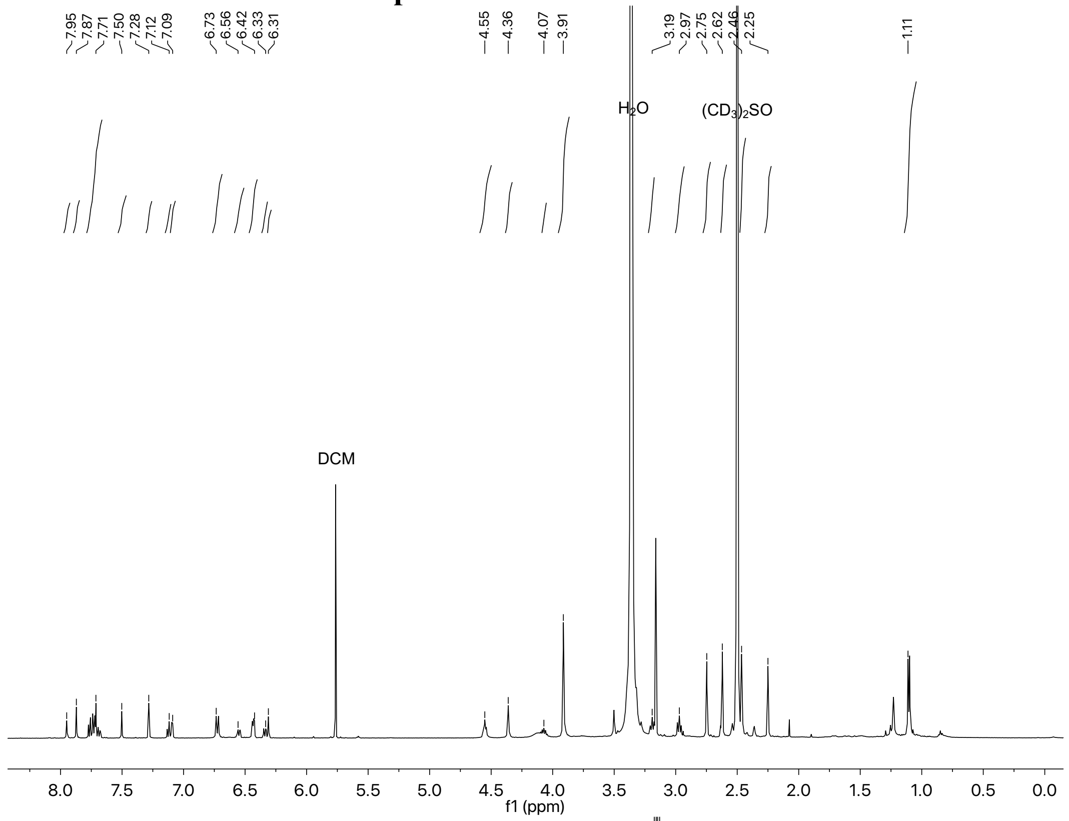

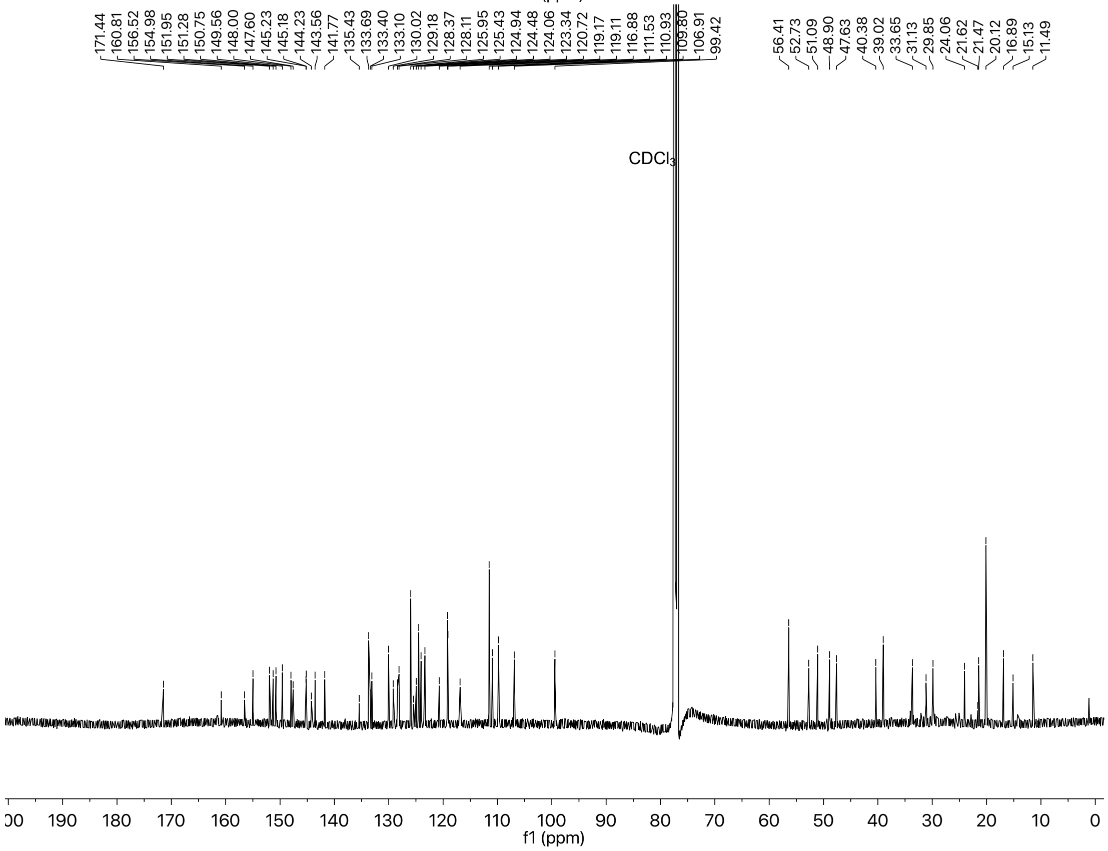

**Compound 3 $^{1}$H and $^{13}$C NMR**

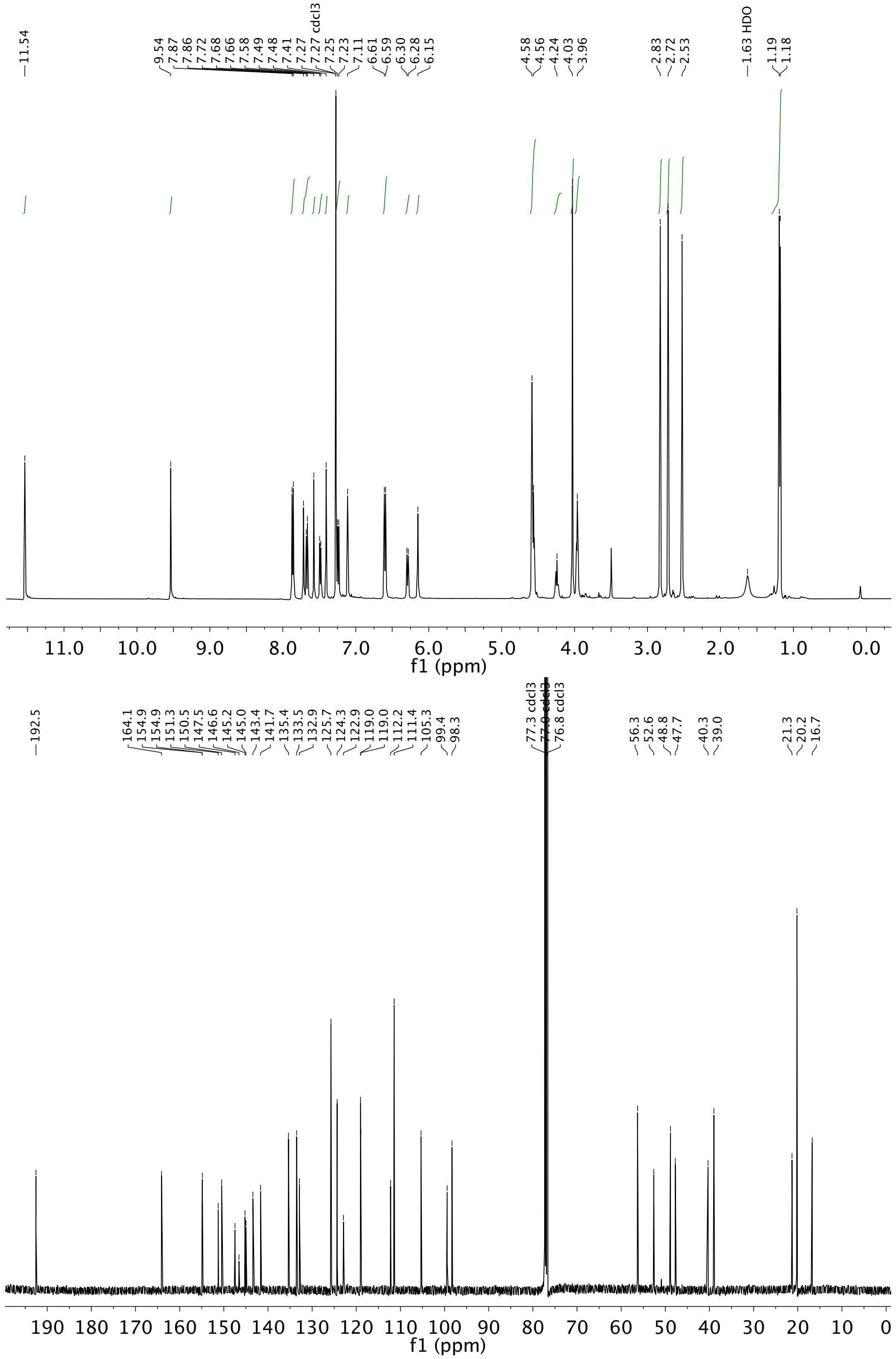

#### Compound 4 $^1$H and $^{13}$C NMR

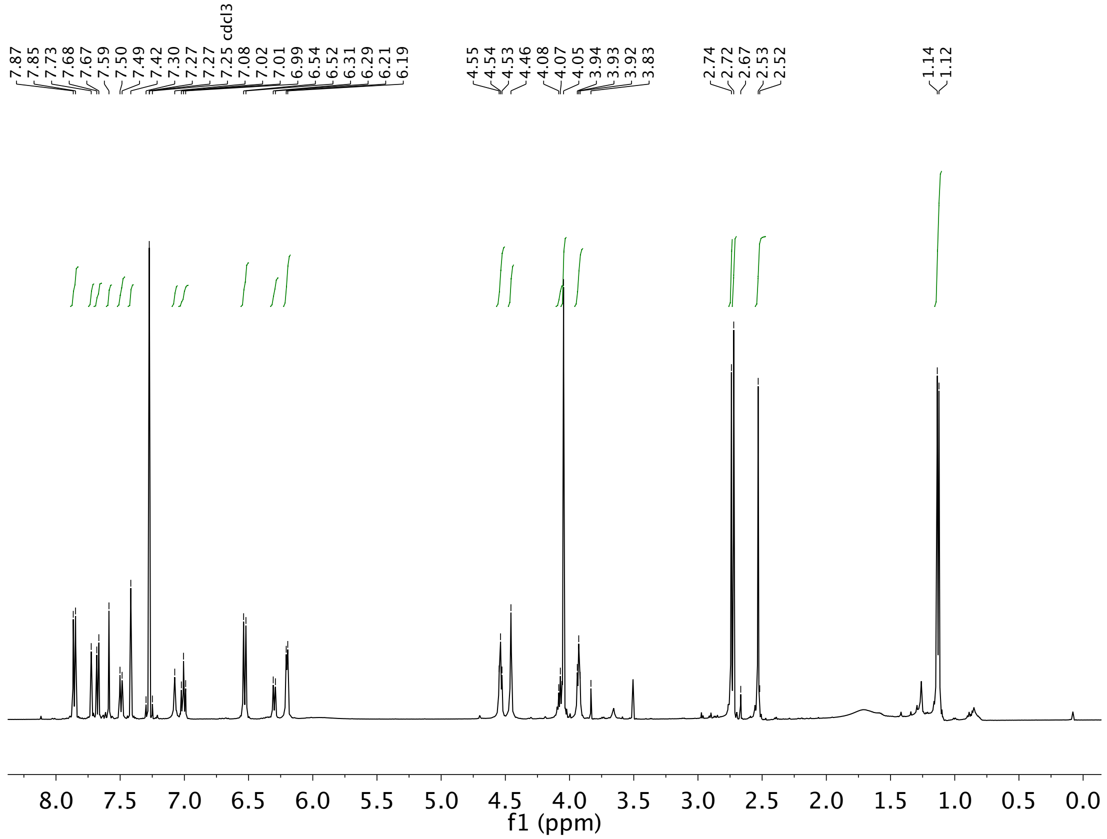


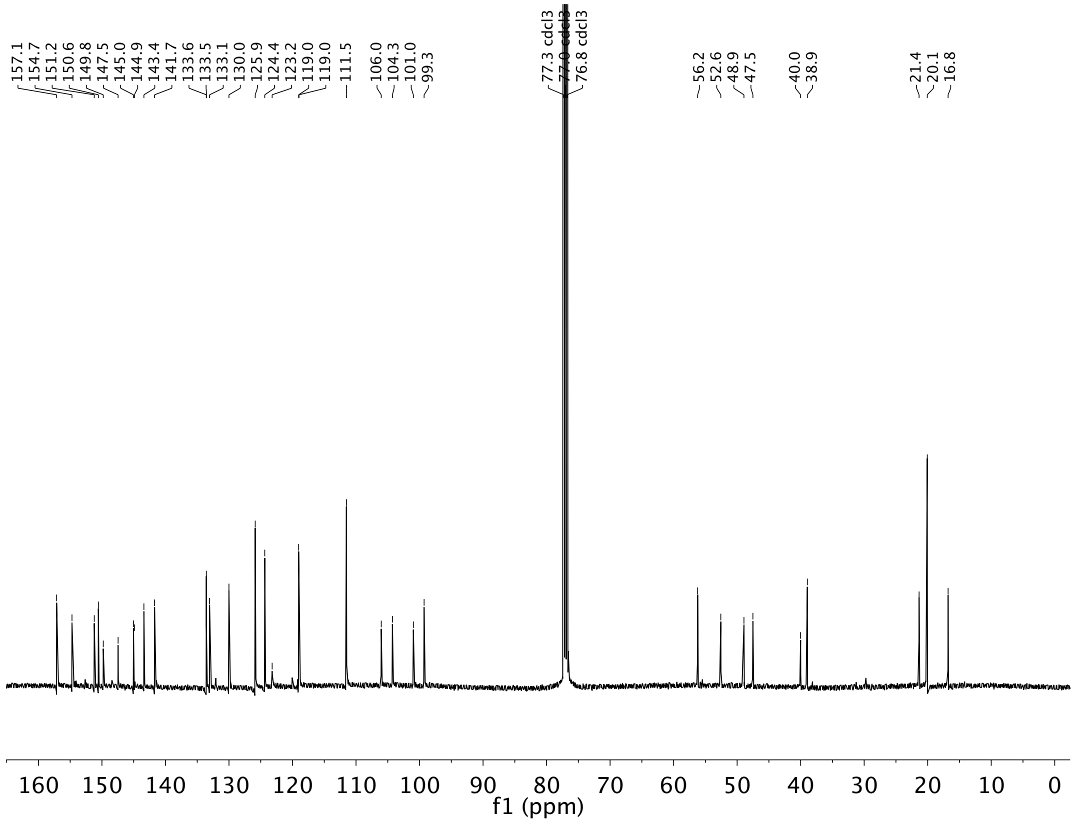

### Compound 5 $^{1}$H and $^{13}$C NMR

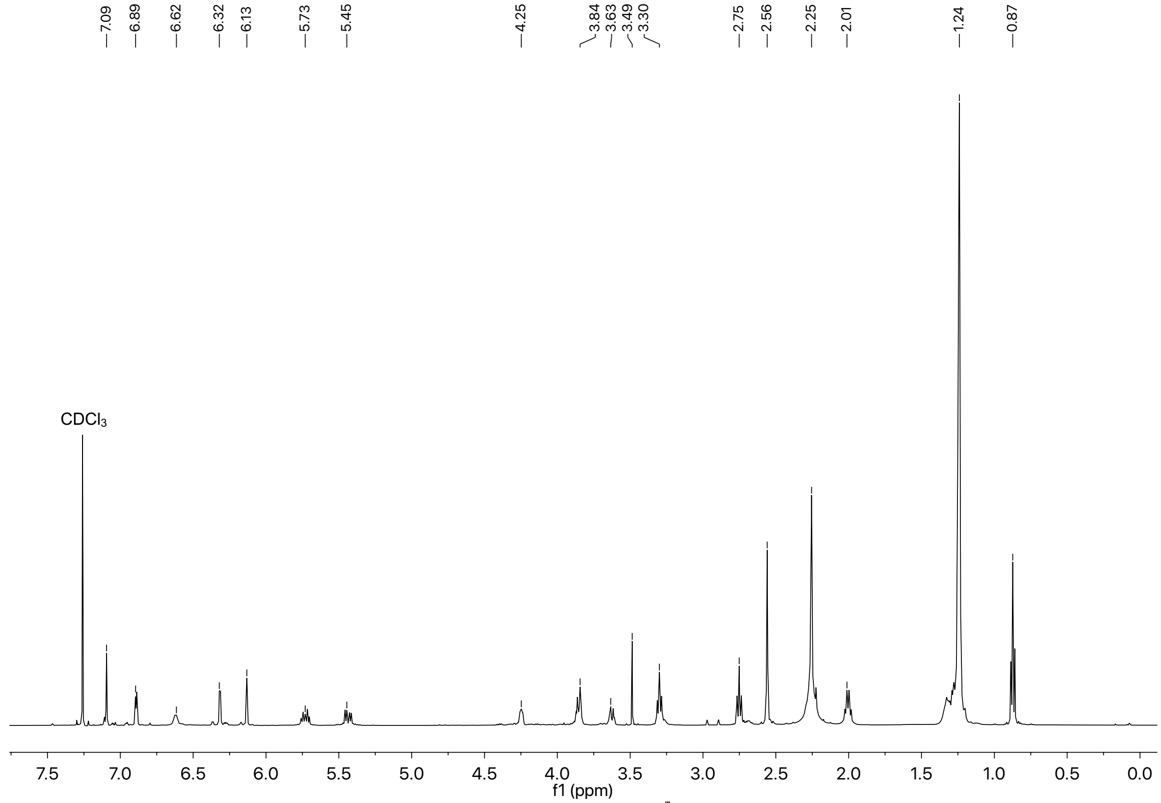


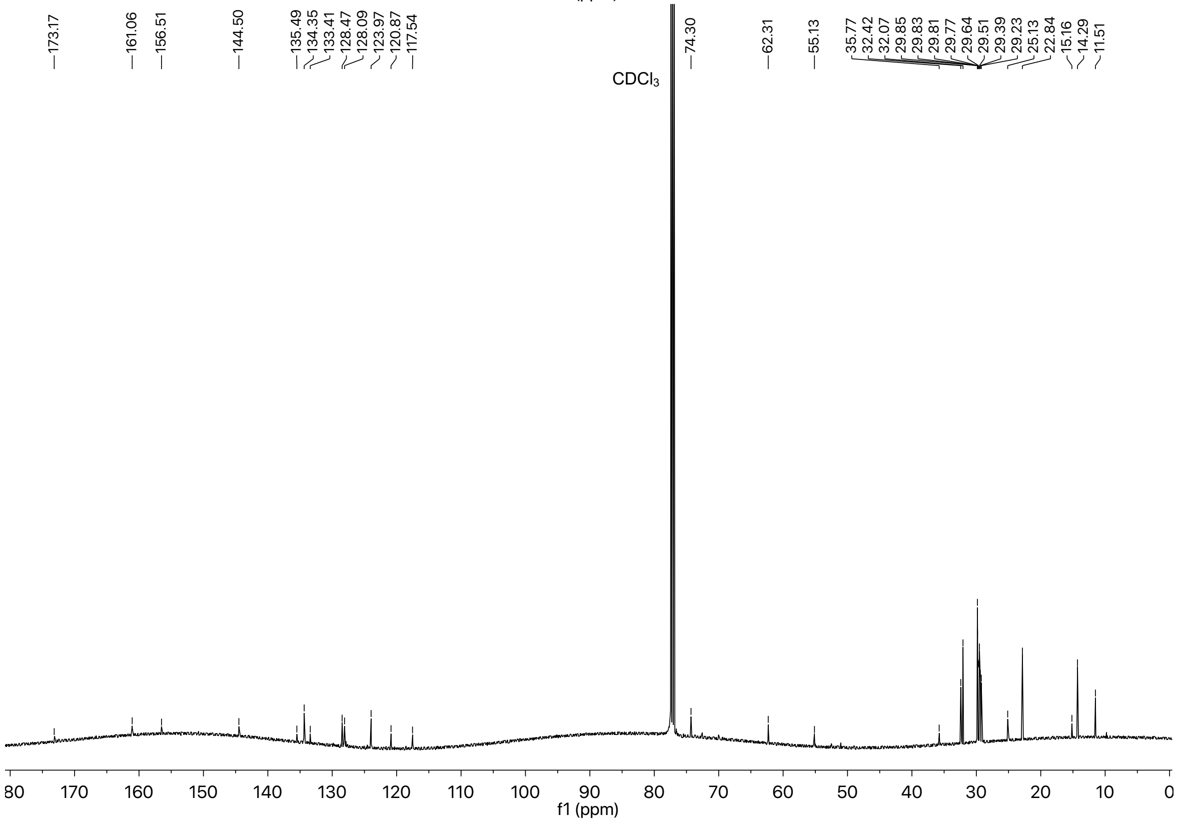

### Compound 6 $^1$H and $^{13}$C NMR

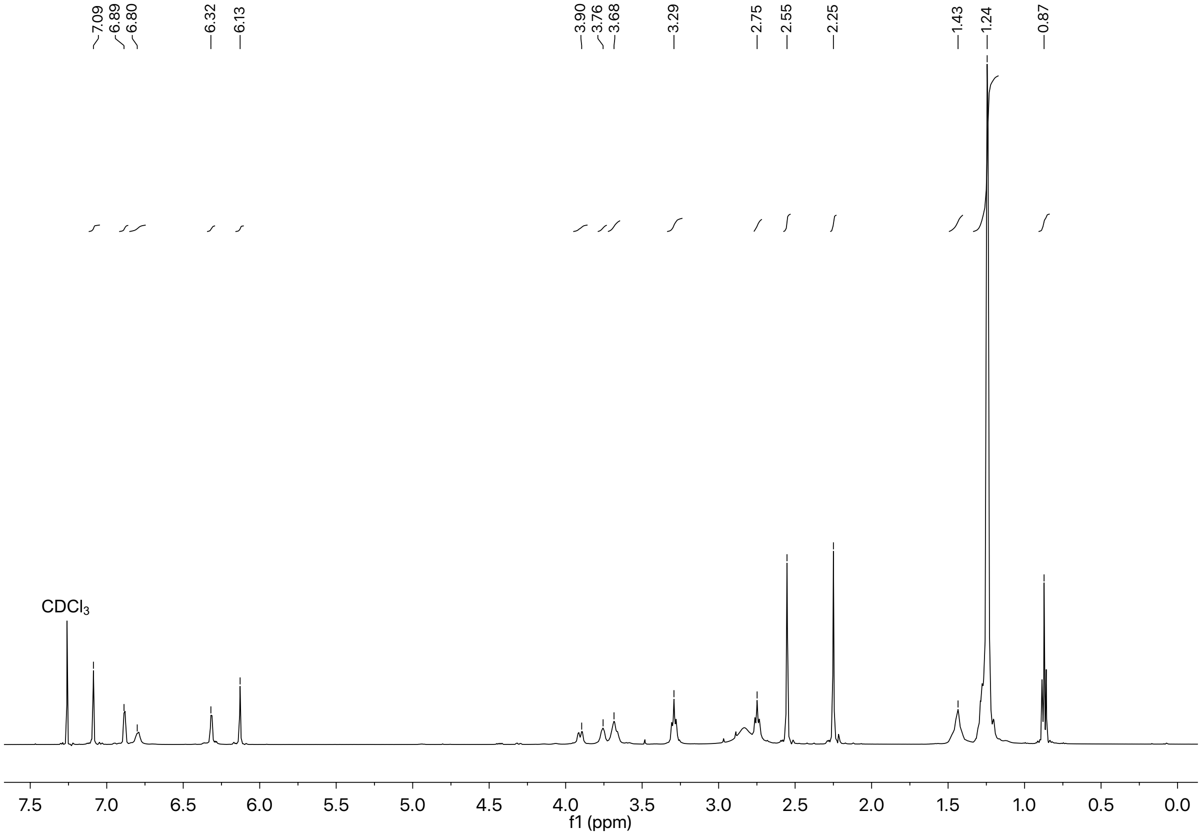


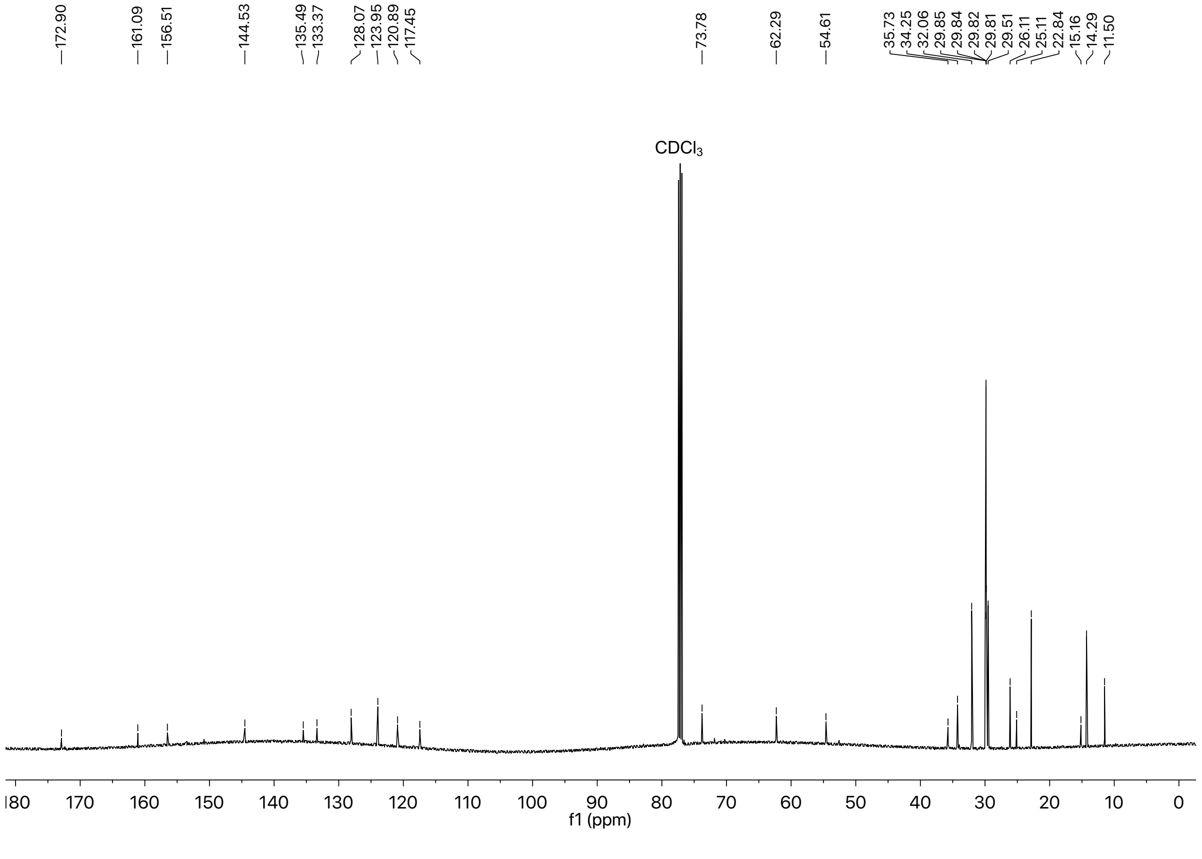

### Mass Spectra

#### Side Product *

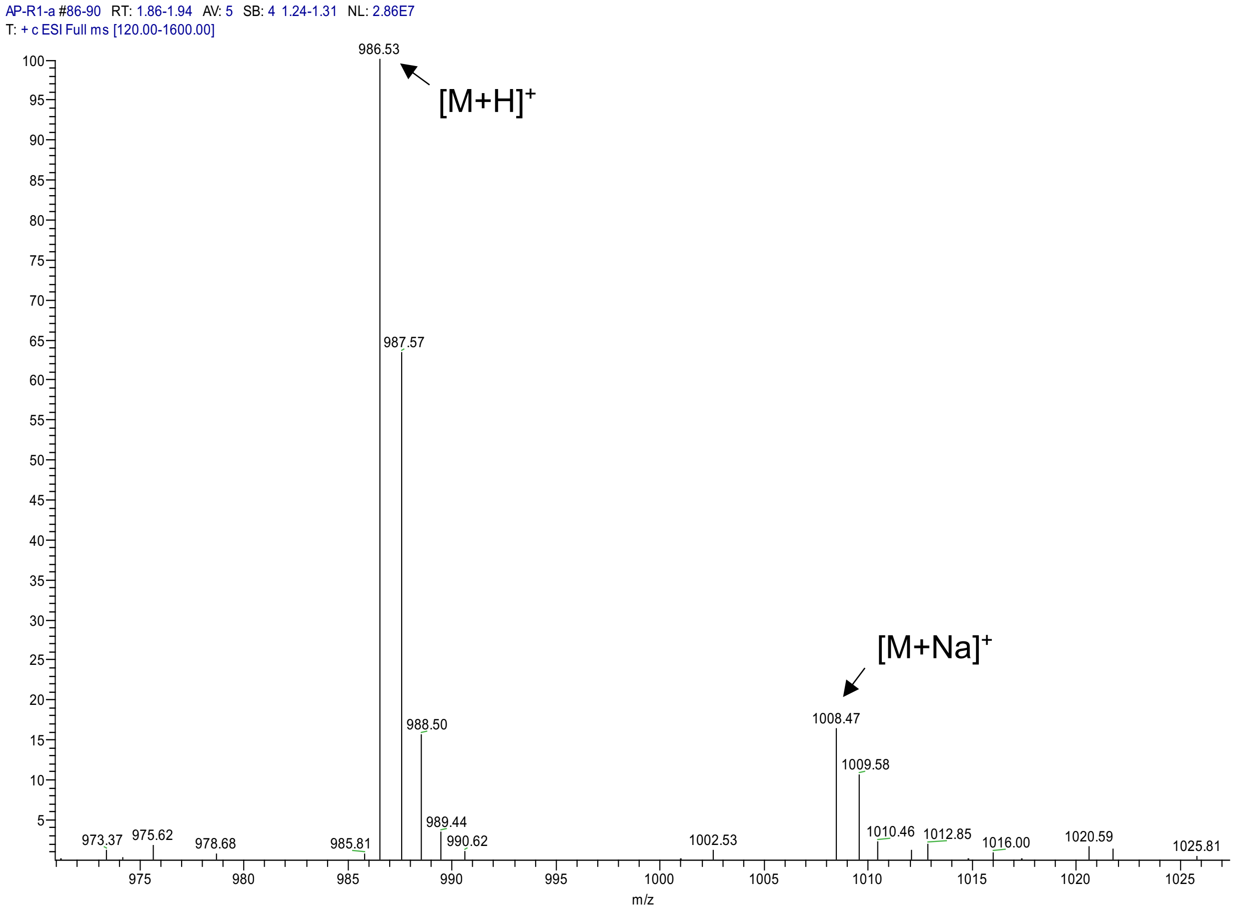

#### Side Product **

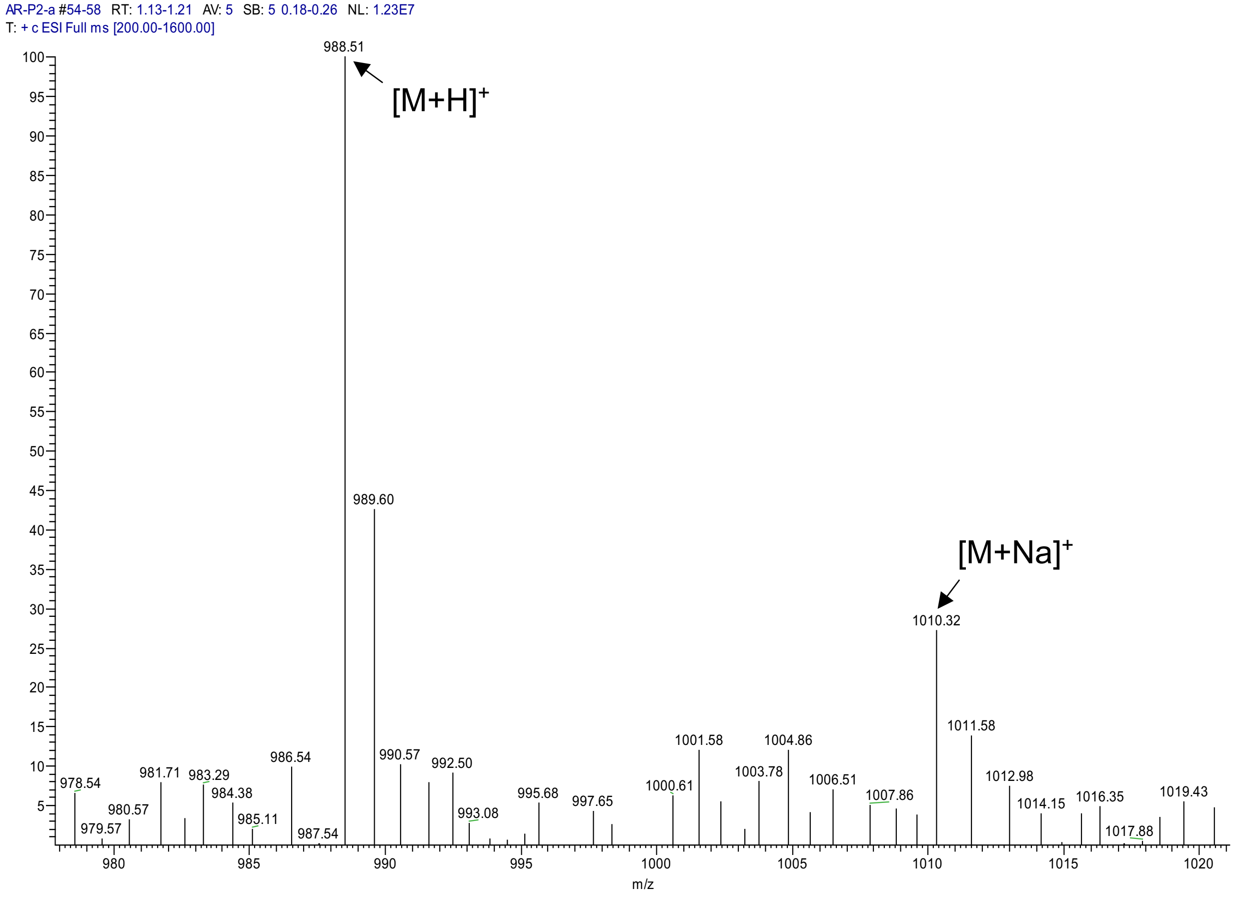
